# Community assembly dynamics in a regulated stream: Insights from environmental DNA

**DOI:** 10.64898/2026.10.11.748123

**Authors:** Ann E. Holmes, Andrew L. Rypel, Mackenzie C. Miner, Dennis E. Cocherell, Tien Ly, Serra C. Perry, Cristina Gonzalez, Nann A. Fangue, Andrea Schreier

## Abstract

Degradation of freshwater habitats, including the alteration of natural flow regimes by impoundment, is a major threat to native fishes. When dam removal is not possible, management actions to recover natural flow regimes can benefit native species, but monitoring community responses is crucial to demonstrating efficacy. We used eDNA metabarcoding to quantify spatiotemporal trends in fish assemblages along the longitudinal profile of a regulated stream where a court-ratified framework uses environmental flows to support native fishes. At a subset of sites, we also evaluated concordance of eDNA results against concurrent surveys using long-term standardized fisheries monitoring techniques. From 59 eDNA samples collected over approximately 2 years, we detected 26 fish taxa, including 8 native species. Twenty-five taxa resolved to species level and one taxon could not be resolved between closely related species. Fish assemblages varied predictably along the longitudinal profile of the stream and by season (fall and spring). Diversity indices were higher at downstream sites and in fall, reflecting greater richness of warm-water non-native species. Evenness was similar among sites but shifts in species ranks suggests reordering processes drive spatiotemporal variation in assembly. Gradient forest analysis identified 13 species associated with spatiotemporal gradients, including both native and non-native species. Overall, eDNA results were concordant with conventional fisheries monitoring, but eDNA detected significantly greater species richness through increased detection of rare species. eDNA results align with long-term monitoring that demonstrated assemblage shifts in only upper stream reaches, and extends the insights into spatial responses, species reordering, and ecology of rare species. These findings demonstrate the value of eDNA as a complementary tool for understanding community assembly dynamics, including responses to stream rehabilitation and ecosystem management.

## Introduction

Freshwater habitats cover only ∼1% of the earth’s surface but are vital habitat for a disproportionate share of global biodiversity (Strayer & Dudgeon, 2010). More than 15,000 described fish species, representing over half of all known ray-finned fishes (Actinopterygii), are found in freshwater habitats (Arthington et al., 2016). Freshwater ecosystems are also an exceptionally valuable natural resource, contributing to nutrient cycling (Vanni, 2002), agriculture and food security (Lund, 2016; Parisek et al., 2024), economic development (Folke, 2003; Gleick & Cooley, 2021), and cultural identity (Reid et al., 2021; Rypel et al., 2021). Yet freshwater fishes face high rates of extinction primarily because of poor ecosystem management including altered flow regimes (Poff et al., 1997; Yarnell et al., 2015), habitat loss and modifications (Sass et al., 2017), and global climate change (Harrison et al., 2018; Heino et al., 2009; Lynch et al., 2023; Strayer & Dudgeon, 2010).

Flow alteration and resulting impacts are widespread. In the northern hemisphere, it is estimated that 77% of large rivers are severely impacted by dams (Dynesius & Nilsson, 1994). In the contiguous United States, there are only 42 remaining high-quality, free-flowing rivers over 200 km in length (Benke, 1990). In lotic ecosystems, flow is a major driver of ecological community structure and function (Lytle & Poff, 2004; Poff et al., 1997); thus, alterations to natural flow regimes negatively impact assemblage diversity (Bunn & Arthington, 2002) and stability (Jacinto et al., 2023). Dams and water diversions drive irreversible change to aquatic habitats and communities (e.g., Moyle, 2014; Pokhrel et al., 2018; da Rocha et al., 2019).

Across the arid to semi-arid North American West, modern dam infrastructure supports water needs including agriculture and flood control, while also driving the decline of native freshwater fish (Brown & Bauer, 2010; Moyle et al., 2011). In California, the use of dams to capture and store water for later use (Lund, 2016) significantly alters hydrologic conditions relevant to native fishes (Null et al., 2014; Zimmerman et al., 2018). While dam removal can help restore native fish populations (Watson et al., 2018) and provide broader social, cultural, and justice benefits (Diver et al., 2024), it is not always feasible or practical (Lejon et al., 2009). For example, certain parts of California with historically high flood risk depend entirely upon dams to protect population centers (Pinter et al., 2019; Thompson 1960).

Nevertheless, human-dominated environments can still support some components of native biodiversity and ecosystem function (Rosenzweig, 2003). For example, flow regimes in highly altered streams can be managed to benefit native freshwater biodiversity (e.g., Jacinto et al., 2023; Konrad et al., 2011; Tonkin et al., 2021). Given the ubiquity of dams, detailed studies on the assembly dynamics of ecological communities in regulated streams are essential, especially in relation to assessing the efficacy of experimental flow regimes (Richter et al., 2003; Yarnell et al., 2020).

Understanding of fish community assembly is limited by the labor- and cost-intensive aspects of conventional sampling (Evans et al., 2017; Ward et al., 2017). The efficacy of conventional fisheries surveys (e.g., electrofishing, traps) can be limited when environmental conditions (e.g., flow) constrain gear deployment or by gear bias due to fish size, species characteristics (Price & Peterson, 2010), or behavioral avoidance (Bouska et al., 2017). Environmental DNA (eDNA) metabarcoding is a non-invasive approach for characterizing biodiversity using trace DNA in environmental samples such as water.

For fisheries monitoring, eDNA methods can be more cost-effective and less labor-intensive (Lafferty, 2024) than conventional methods and can reduce or eliminate the potential for fish mortality and stress during sample collection. In freshwater systems with fewer than 100 fish species, eDNA metabarcoding detects more species than conventional surveys due to a higher sensitivity for rare species (McElroy et al., 2020). Further, eDNA is effective for detecting endangered species and invasive species, including previously unrecorded invasive fishes (Feng et al., 2023). For the dominant species in fish communities, eDNA metabarcoding results reflect relative abundance (Skelton et al., 2023), enabling detection of changes in fish community structure that may be missed by conventional surveys.

eDNA metabarcoding captures management-relevant spatiotemporal patterns and species turnover in aquatic invertebrate communities with higher resolution than conventional sampling (Seymour et al., 2021), but similar analyses using eDNA have yet to be widely applied to fish communities. In this study, we: (1) use eDNA metabarcoding to elucidate spatiotemporal trends in fish communities across an environmental gradient and between seasons, (2) compare findings with conventional fisheries monitoring using multiple gear types by leveraging opportunistic overlap of these standardized surveys with eDNA sampling, and (3) evaluate the utility of eDNA for informing science-based conservation and management of native fishes in a regulated stream.

## Methods

### Study site

Lower Putah Creek is a moderately sized stream of ∼40 km (25 mi) in the Central Valley of California, USA (Figure 1). The name ‘Putah’ may derive from the local Indigenous placename *puta-to*, a historic Patwin village in the Berryessa Valley (Gudde, 2004). Originating in the Northern Coast Range, Putah Creek flows eastward ∼130 km (80 mi) before reaching Lake Berryessa, a reservoir (capacity = 1.98 km^3^) regulated by Monticello Dam (completed in 1963). About 10 km (6 mi) downstream of the dam, a secondary structure, the Putah Diversion Dam (PDD), diverts water south for domestic and agricultural uses. Below the PDD, an environmental gradient is driven by upstream water releases and downstream warming (Kiernan et al., 2012; Jacinto et al., 2023). The creek terminates at the Yolo Bypass, a managed floodplain in the Sacramento River delta and part of the San Francisco Estuary, the largest estuary on the Pacific coast of the United States.

**Figure 1.**
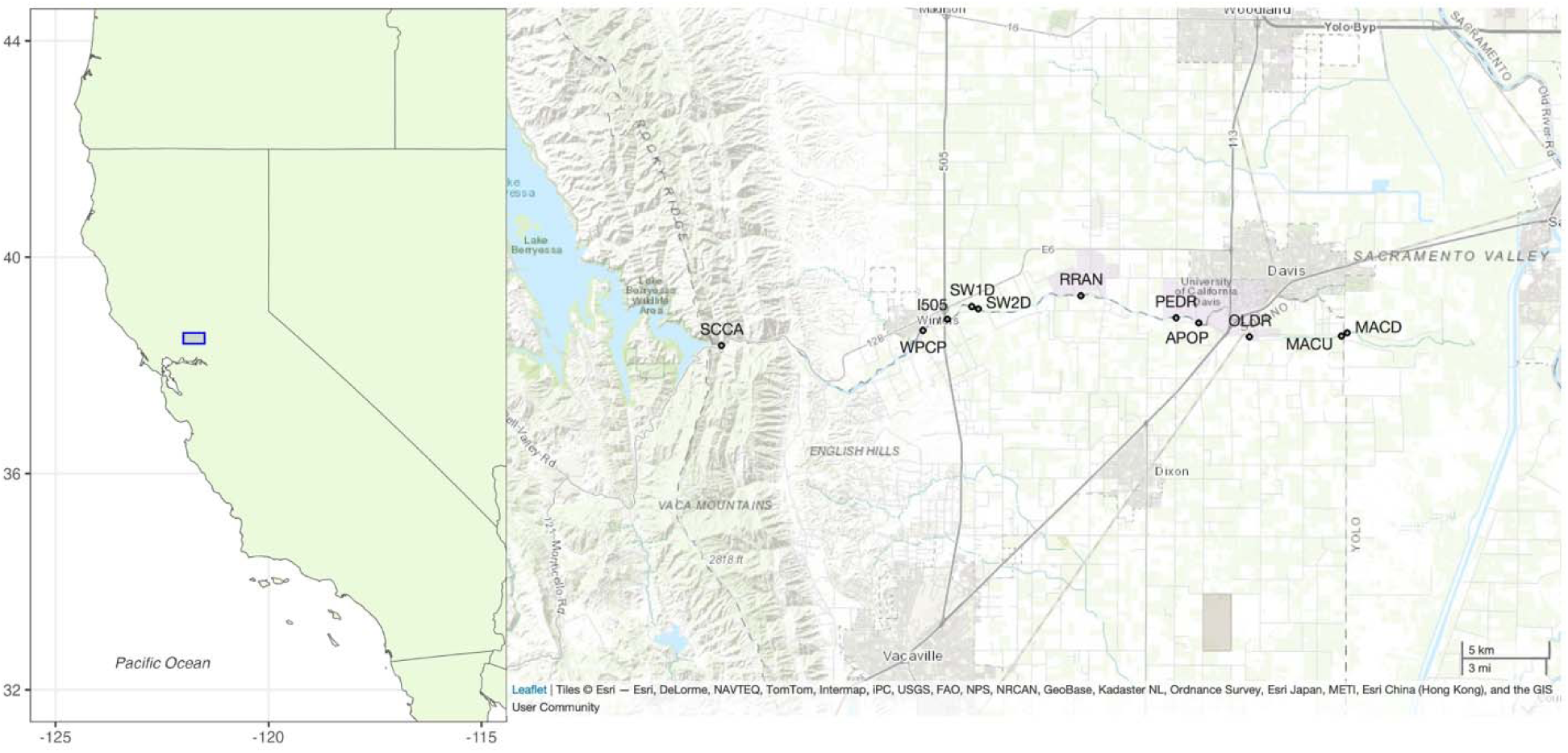
Map of lower Putah Creek (Solano and Yolo Counties, California) showing eDNA sampling sites (2019-2021); site names corresponding to map labels can be found in Table 1.

**Table 1.** Sites in Lower Putah Creek (Solano and Yolo Counties, California) sampled for environmental DNA (eDNA) from 2019-2021. A subset of eDNA sites was also sampled at least once with a concurrent (or within 24 hours) conventional survey as shown. “Multiple” methods represent a comprehensive sampling effort including electrofishing, traps, gill nets, and beach seine.

| Site name | Site code | Distance from Yolo Bypass confluence (km) | Latitude | Longitude | Concurrent conventional survey |
| --- | --- | --- | --- | --- | --- |
| Mace Boulevard, downstream | MACU | 10.6 | 38.5192 | -121.6967 | none |
| Mace Boulevard, upstream | MACD | 11.1 | 38.5176 | -121.6967 | electrofishing (2020, 2021) |
| Old Davis Road | OLDR | 16.9 | 38.5172 | -121.7562 | electrofishing (2020); none (2021) |
| Fire Ring Picnic Grounds | FRPG | 19.2 | 38.5242 | -121.7889 | multiple (2021) |
| Pedrick Road | PEDR | 20.8 | 38.5268 | -121.8035 | electrofishing (2020, 2021) |
| Russell Ranch | RRAF | 26.7 | 38.5379 | -121.8650 | fyke trap (2020) |
| Step Weir 2 | SW2D | 34.1 | 38.5313 | -121.9313 | rotary screw trap (2021) |
| Step Weir 1 | SW1D | 34.4 | 38.5325 | -121.9355 | none |
| Interstate 505 Crossing | I505 | 35.1 | 38.5261 | -121.9513 | electrofishing (2021) |
| Winters Putah Creek Park | WPCP | 36.5 | 38.5204 | -121.9670 | electrofishing (2021) |
| Stebbins Cold Canyon | SCCA | 51.0 | 38.5128 | -122.0972 | none |

Putah Creek is habitat for formerly abundant native freshwater and anadromous fishes (Moyle et al., 1998). The creek historically experienced high seasonal and annual variations in streamflow typical of Mediterranean-type climates. Summers are hot and dry; most rainfall occurs during the winter, often resulting in high-flow events and seasonal inundation. Following decades of degradation caused by dams and diversions, a legal settlement in the 1990s resulted in the Putah Creek Accord (Börk et al., 2012; Moyle et al., 1998), initiating a management strategy based on the natural flow regime concept (Poff et al., 1997). Since its implementation, data from annual, long-term electrofishing surveys suggest native fish communities have benefitted from the restored flow regime (Kiernan et al., 2012; Jacinto et al., 2023; Hitt et al., 2025).

### eDNA sample collection

eDNA methods followed rigorous protocols to minimize contamination risk in the field and laboratory (Goldberg et al., 2016). Prior to field work and sampling, equipment was sterilized with 20% bleach solution for 20 minutes, then rinsed twice with deionized water. All laboratory work was conducted in spaces free of fish, fish tissue, and fish DNA. One field negative control sample was included for each sampling day (maximum 3 sites sampled per day) to assess potential contamination from handling, cross-contamination, equipment, or the environment; a 1-L bottle of deionized water was transported with field gear, opened at the field site for approximately 1 min, and thereafter filtered, prepared, and sequenced alongside the field samples.

eDNA field samples were collected in 21 sampling events from 12 sites over ∼2 years (December 2019 to October 2021; Figure 1; Table 1; Table S1). Sites were selected for spatial coverage along environmental gradient of Putah Creek and to prioritize opportunistic concurrent sampling with long-term fisheries surveys described below. Each site was sampled 1 to 3 times, with variation in sampling effort due to impacts of the COVID-19 pandemic and land access restrictions. Samples were collected in either fall (October to December) or spring (April to May). Replicate water samples (n=3) were collected from the bank in sterile 1-L bottles using a swing sampler (Nasco, Atkinson, Wisconsin, USA). For collections at Step Weir 2 (SW2D; see Table 1), water sampling was conducted from a small boat in the center of the creek used to access the rotary screw trap.

Water was filtered using 47 mm diameter 1.6 µm pore glass fiber filters (Whatman International Ltd., Maidstone, UK) and an eDNA sampler (Smith-Root, Inc., Vancouver, WA, USA) or a peristaltic pump (Geotech Environmental Equipment, Inc., Sacramento, CA, USA) running at a rate of 10-15 minutes per liter. Filters were folded in half twice, placed in 2 mL microcentrifuge tubes, and frozen at -20°C until DNA extraction, which was typically done within 4 weeks. Negative control samples were filtered and preserved in the same manner.

### Conventional fish surveys

Except for the most upstream eDNA sampling site (Stebbins Cold Creek; Figure 1), historical occurrence data is available via long-term fisheries monitoring (Jacinto et al., 2023). In addition, 12 of the 21 eDNA sampling events are matched with concurrent conventional fisheries monitoring (Table 1). Only a subset of eDNA samples have corresponding conventional data because concurrent conventional monitoring was conducted independently and the gear used depended on the season. Fall electrofishing, part of long-term fisheries monitoring, was the most spatially extensive. The other surveys (rotary screw trap, fyke trap, and multi-method) each occurred at a single site.

Electrofishing surveys used a tote barge electrofishing (or rarely backpack electrofishing, if the creek was inaccessible to the barge) consisting of single-pass, equivalent stream-length distances conducted annually in October following established protocols (Jacinto et al., 2023).

These surveys have occurred annually since 1996. Stunned fish were collected by net and enumerated, identified, measured, and released. Water for eDNA analysis was collected less than 24 hours prior to conventional sampling, usually on the same day.

Rotary screw trap surveys, the standardized method for monitoring out-migrating juvenile salmon, were conducted following established protocols (Rypel et al., 2022; Miner, 2022). In 2020, fyke nets were deployed instead because installation and operation of the rotary screw trap was not feasible during the COVID-19 pandemic for staffing and safety reasons.

Spring rotary screw trap or fyke sampling has occurred annually in the spring since 2017. Water for eDNA analysis was collected just downstream of traps approximately 10-30 min before traps was opened for fish identification, enumeration, and measurement.

The multi-method survey, which combined electrofishing, fyke traps, and seining, was conducted by instructors and students as part of a university fish biology course. These surveys have occurred annually since the mid 1970s. Water samples were collected at the midpoint of the survey area as survey teams deployed.

### DNA extraction, amplification, and sequencing

DNA was extracted from whole filters using the PowerWater DNeasy Kit (Qiagen, Hilden, Germany) following the manufacturer’s instructions with minor modifications (Holmes et al., 2024). Extracted DNA was stored in LoBind tubes (Eppendorf, Hamburg, Germany) at - 20° C. eDNA metabarcoding library preparation followed a two-step PCR protocol (Illumina, 2013). The first (amplicon) PCR used the MiFish-U amplicon primer set targeting a ∼170 bp portion of the fish 12S ribosomal RNA (rRNA) gene (Miya et al., 2015) and the KAPA HiFi PCR kit (Roche Diagnostics, Indianapolis, USA), with an annealing temperature of 68°C and 35 cycles; details are provided in Supplementary Tables S2–S4. PCR conditions and cycling protocol for the second (index) PCR, which attaches unique sample-specific indices and sequencing adaptors, followed Illumina (2013) guidelines. Sample libraries were pooled in equimolar amounts, checked for correct amplicon size using a Bioanalyzer and sequenced on an Illumina MiSeq (2x150 paired-end reads) at the University of California-Davis DNA Technologies and Expression Analysis Core (Davis, CA) with 15% Phi-X added.

In addition to the field negative controls, extraction and laboratory controls (no-template PCR reactions) were included at each step. All negative controls were checked for contamination using gel electrophoreses. A small volume (1 µl) of nine field negative controls and one laboratory control were added to the pooled library as an additional quality control measure.

### Bioinformatics and taxonomic assignment

Bioinformatic analysis and data processing were performed in R version 4.2.1 using cutadapt v3.7 (Martin 2011), DADA2 v1.25.0 (Callahan et al., 2016) and phyloseq v1.44.0 (McMurdie & Holmes, 2013) packages. First, cutadapt was implemented in the R environment to trim amplicon primers from demultiplexed sequence reads. Reads were filtered and trimmed using DADA2’s `filterAndTrim()` function using error rates calculated from sequencing quality scores and retaining sequences 150–195 bp. Sequences >195 bp are typically off-target amplification of prokaryote taxa, a known artifact of the MiFish primer set (Miya et al., 2015). Amplicon sequence variants (ASVs) were inferred and denoised in using modeled error rates. ASVs are exact, error-corrected sequences that represent unique variants of the MiFish barcode. A pseudo-pooling approach was used to help retain rare ASVs across the dataset while removing errors. Forward and reverse reads were merged, and chimeric sequences were removed.

Taxonomy was assigned to ASVs using DADA2’s `assignTaxonomy()` function, which implements the Ribosomal Database Project (RDP) Naive Bayesian classifier algorithm (Wang et al., 2007), and a custom regional reference sequence database modified from Nagarajan et al. (2023). The database included all fish species detected in long-term fishery monitoring of Putah Creek (Jacinto et al., 2023) and other fishes know from the upper San Francisco Estuary. Aquatic, semi-aquatic and riparian vertebrates (i.e., turtles, mammals, and birds) and potential trace contaminant DNA, including human and taxa from concurrent projects (e.g., rockfish (*Sebastes* spp.), a marine taxon) were also included in the reference sequence database. When barcode sequences were identical between related species, the taxa were annotated to genus or family level and the sequences dereplicated. Taxonomic assignment was tested using bootstrap values of 0.5 and 0.80. The default bootstrap parameter in DADA2 is 0.5, but values of 0.7-0.8 are common in taxonomic assignment (Wang et al., 2007).

The ASV table was decontaminated by filtering ASVs with <100 reads and non-fish ASVs from the dataset. To further reduce risk of false positive detections due to trace laboratory contamination and artifacts of amplicon sequencing (i.e., “tag-jumping;” Schnell et al., 2015), the fish ASV table was decontaminated by subtracting, from each sample, the maximum number of sequence reads observed of taxa detected in any negative control sample. Taxonomic assignments were also checked for concordance with species reported in historical surveys (Jacinto et al., 2023). Unexpected species detections and unassigned ASVs were cross-checked against NCBI GenBank’s core nucleotide database via manual BLAST using default settings (Altschul et al., 1990). ASVs were agglomerated to species level using phyloseq’s `taxa_glom()` function. Subsequent data analysis and visualization was conducted at the species level. Sequencing results were visualized using species accumulation curves to assess whether sequencing depth was sufficient to detect species diversity at each sampling event.

### Community composition and spatiotemporal analyses

Site-level occurrence of fish species was visualized in a phylogenetic framework based on MiFish reference sequences. Sequences were aligned using the `AlignSeqs()` function in the R package DECIPHER v3.8.1(Wright, 2024). The `dist.dna()` function in the R package ape v5.8.1 (Paradis & Schliep, 2019) was used to calculate pairwise genetic distances using p-distance (proportion distance) and pairwise deletion to account for missing or ambiguous bases; species pairs with genetic distances <3% were flagged for further consideration. A UPGMA tree was constructed using the resulting distance matrix with Pacific lamprey (*Entosphenus tridentatus*) set as the outgroup and visualized using ggtree v4.2.0 (Yu et al., 2017). The tree and genetic distances are presented to help evaluate reliability of species assignments; short barcode sequences do not necessarily reflect true evolutionary relationships.

Community and spatiotemporal analyses were conducted using the R package vegan (Oksanen et al., 2022). Relative read abundance of fish species was calculated for each sample. Species richness, Shannon diversity index, and evenness (Pielou’s J) were calculated using mean relative read abundance for all replicates in a sampling event. For community assemblage (beta diversity) analysis, eDNA detection data was converted to eDNA Index following Kelly et al., (2019). eDNA Index values are the relative abundance of each species normalized across samples and account for differential amplification of species in eDNA metabarcoding (e.g., primer bias; Kelly et al., 2019), allowing for more meaningful comparisons in multivariate analyses. Community dissimilarity was visualized using non-parametric multidimensional scaling (NMDS) based on Bray-Curtis distances. A stress value was calculated to determine the ordination’s goodness-of-fit. Permutational multi-variate analysis of variance (PERMANOVA), also using eDNA Index values, was used to determine significance of compositional dissimilarity by sampling site and season using the adonis2 function based on Bray-Curtis distances in the vegan package.

### Comparison with conventional sampling

eDNA results were compared with concurrent electrofishing and trap surveys using alluvial plots created with the R package ggalluvial (Brunson & Read, 2023). For both electrofishing and trap samples, individual counts by species were compared with eDNA relative read abundance. For trap samples, biomass data was also available and compared with eDNA relative read abundance by species. Species richness of eDNA samples was compared with concurrent electrofishing surveys (n=7) using a Wilcoxon signed-rank test (α = 0.05) with a one-tailed hypothesis. Statistical comparisons were not made with trap samples or the multi-method survey due to a small sample size (1-2 paired sampling events per year).

### Reordering and species-specific patterns

Spatiotemporal reordering in species composition was investigated using rank-abundance curves (also known as dominance-diversity ‘Whittaker plots’) created using the R package codyn (Hallett et al., 2016; Hallett et al., 2020). Species rank was based on relative sequence read abundance and highlighted patterns by native/non-native status and common species. In an exploratory analysis, temporal beta diversity was investigated using two metrics: mean rank shift (MRS) and species turnover. MRS is the mean change in ranks across species and turnover captures species lost or gained. Rank shifts can demonstrate re-ordering of the same species or community change in long-term studies, while turnover captures compositional change across time (Hallett et al., 2016). Higher MRS indicate variable and less stable assemblages, while lower MRS values suggest stable ecological community structure. Annual MRS trends in Putah Creek were previously evaluated based on long-term electrofishing data (Jacinto et al., 2023).

Since limited temporal data were available across sequential years or seasons in our study, these metrics were interpreted with caution. Season-to-season comparisons are unconventional but may provide novel insight into community-level dynamics of Mediterranean-type ecosystems with high seasonality, and potential community ecology applications using eDNA data.

Species-specific responses and variable importance were investigated using a gradient forest model, a machine learning approach that models individual species occurrence to identify those contributing most to assemblage-level change. The model was implemented in the gradientForest package (version 0.1–17; Ellis et al., 2012) using 500 runs with variables of sampling site (river km), date, and year as variables. Parallel to the community ecology metrics, this model was based on eDNA Index values as a proxy for relative species abundance (Kelly et al., 2019). Species with *R^2^*> 0.50 were considered to have strong responses to the variables because a drop off in *R^2^* was identified below 0.50 (broken stick method; Frontier, 1976).

R scripts and the custom reference sequence database are available on Zenodo (add link).

## Results

Metabarcoding of 59 eDNA field samples produced ∼4.9 M raw sequence reads and ∼3.1 M quality-filtered fish sequence reads (mean 52,284 reads/sample; Supplementary Table S5). Species accumulation curves indicated sequencing depth was adequate to capture the full extent of fish species richness in each replicate sample (Supplementary Figure S1). A total of 68 fish ASVs (defined in Methods) passed quality control and were assigned to 26 fish taxa; 25 taxonomic assignments resolved to species level (Table 2; Figure 2). Seven species pairs had genetic distances <3% (Supplementary Table S6). Sequences assigned to non-native black bullhead (*Ameiurus melas*) and non-native brown bullhead (*A. nebulosus*) were combined because the species had identical MiFish barcodes (Nagarajan et al., 2023). In addition, non-native silversides (*Menidia audens* and *M. beryllina)* were treated as a species complex; genetic data from silversides indicates morphological divergence may reflect rapid adaptation to new habitats, rather than a phylogenetic split between two nominal species; thus, taxonomic validity remains uncertain (Fluker et al., 2011). Following Jacinto et al. (2023), all sculpins were identified as *Cottus asper*.

**Figure 2.**
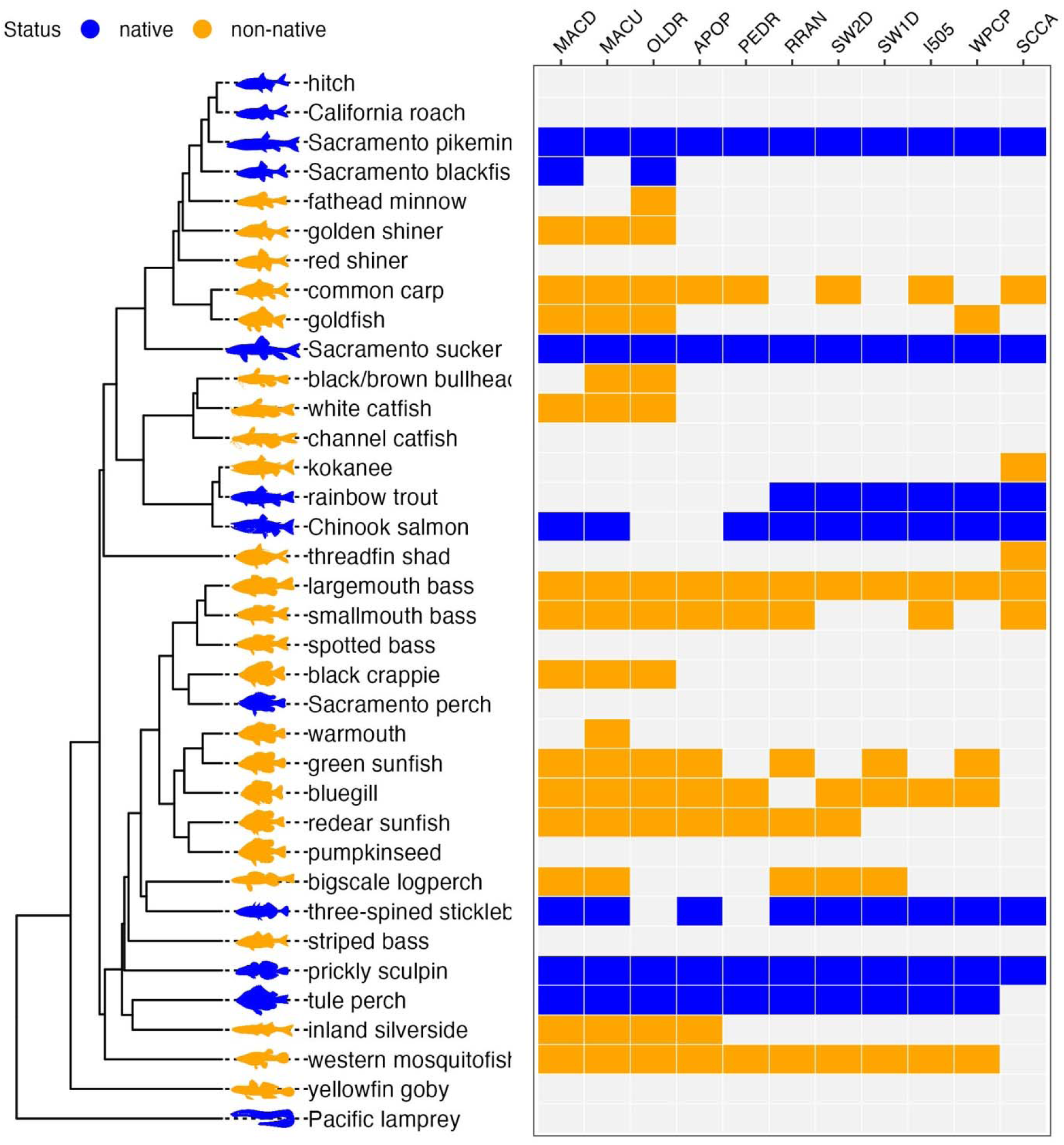
MiFish barcode phylogeny of Putah Creek fishes and site-level eDNA detections. Stations ordered downstream to upstream (left to right) and station codes can be found in Table 1. Fish taxonomy is provided in Table 2. Details on the fish silhouette source files and licensing information are provided in the Zenodo repository.

**Table 2.**
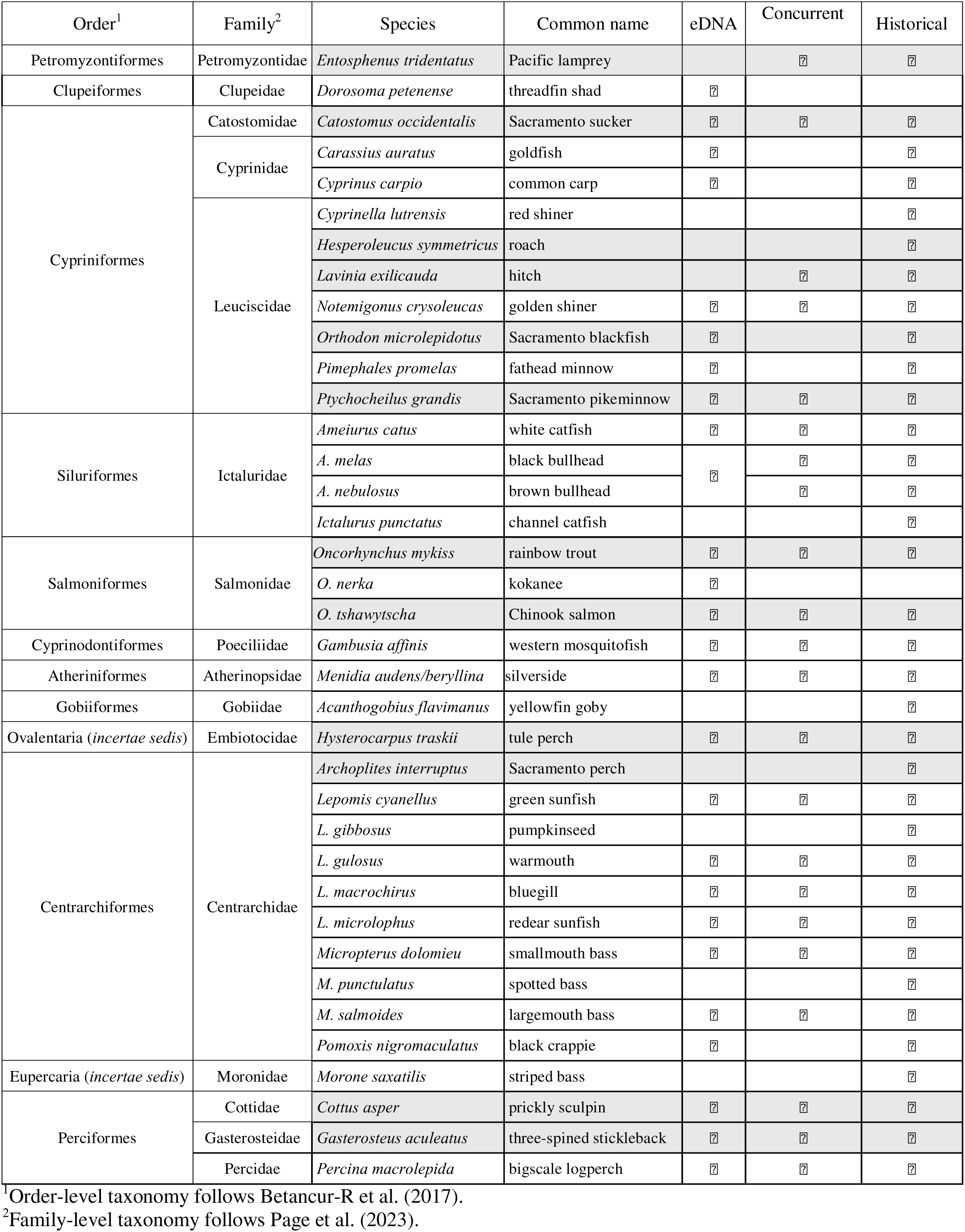
Fish species detection using eDNA metabarcoding and conventional sampling conducted concurrently at the same sites (2019-2021) compared with historical data from standardized surveys (1993-2017; Jacinto et al., 2023); native species are indicated by shaded boxes. Threadfin shad and kokanee detections may reflect downstream eDNA transport.

Field and laboratory negative controls showed no evidence of contamination, generating from zero to 635 fish sequence reads (mean 89 reads/sample). A small number of reads (maximum 88) from abundant taxa were subtracted from ASV sequence read counts based on detection of those ASVs in the field negative control samples. These detections are consistent with low-level laboratory contamination or tag-jumping common in Illumina workflows (Hänfling et al., 2016; Schnell et al., 2015) addressed through the decontamination protocol.

ASVs identified as probable laboratory contaminants included marine and mock community species from concurrent projects, suspected to be the result of minor cross-contamination, were also detected in very low abundance and filtered from the dataset. In addition, an ASV that occurred in a single field sample was initially assigned to a regionally occurring species not known to occur in Putah Creek but was reclassified as Atlantic menhaden (*Brevoortia tyrannus*; 100% similarity), a marine species, after validation against NCBI GenBank via BLAST (Altschul et al., 1990). This detection was attributed to low-level contamination from animal feed unexpectedly introduced into a shared, fish-free laboratory where eDNA field gear was stored.

### Community composition and spatiotemporal analyses

Twenty-six fish taxa from 12 families were detected using eDNA, comprised of 8 native species and at least 18 non-native species (Table 2; Figure 2). Species detections were broadly congruent with historical Putah Creek surveys (Jacinto et al., 2023). Nearly 30% of the total sequence reads were identified as native Sacramento sucker or native Sacramento pikeminnow, 15.0% and 14.5%, respectively. Both species were detected at least once at each site (Figure 2). Two species detected by eDNA, non-native kokanee salmon and threadfin shad, have not been reported from Putah Creek but are known from Lake Berryessa upstream. These species were detected in multiple replicates at the most upstream eDNA sampling location (Stebbins Cold Canyon, 51.0 km above the confluence with the Yolo Bypass), just below the reservoir; this site is not included in historical surveys. These detections likely reflect transport of eDNA or fish from the reservoir; kokanee salmon are stocked annually in Lake Berryessa and threadfin shad are occasionally reported the creek, but only when the emergency spillway on Monticello Dam activates. Two ASVs were assigned to prickly sculpin (*Cottus asper*); neither sequence matched inland riffle sculpin (*C. gulosus*), the other freshwater sculpin known from the region.

Non-metric multidimensional scaling (NMDS) of beta diversity (Figure 3) revealed clustering of samples by stream region. The stress value (0.13) indicates an acceptable goodness-of-fit for the ordination. Downstream sites from Mace Road to Old Davis Road (10.6 to 16.9 km from the Yolo Bypass confluence) clustered together and upstream sites from Step Weir 1 to Putah Creek Winters Park (34.1 to 36.5 km from confluence) clustered together. PERMANOVA analysis showed significant community dissimilarity by site (p<0.001) and season (p<0.05). As seen in historical electrofishing data (Jacinto et al., 2023), species richness and Shannon diversity index were higher at downstream sites, while no spatiotemporal trends were apparent in evenness (Supplementary Table S7).

**Figure 3.**
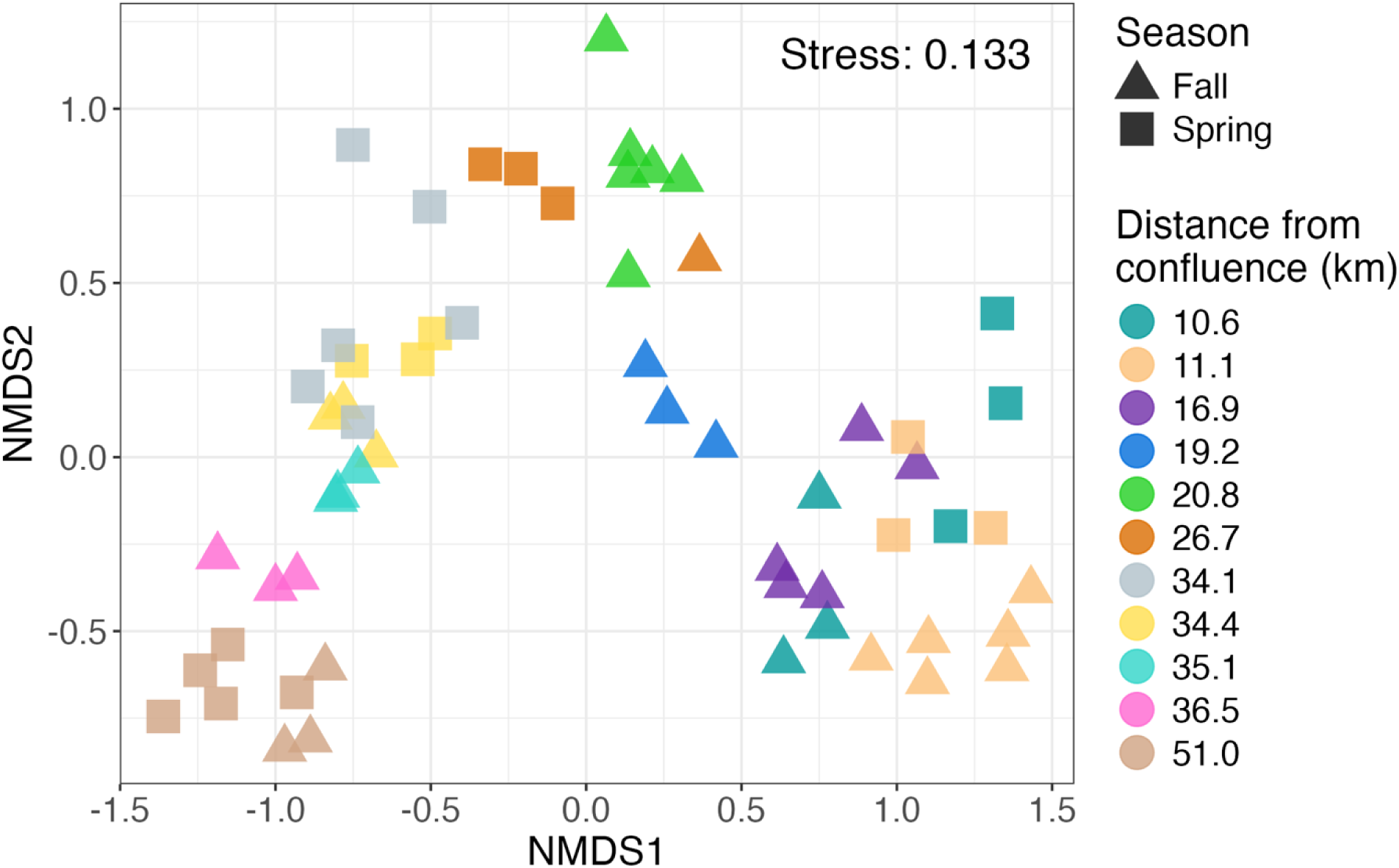
Non-metric multidimensional scaling (NMDS) plot of sample beta diversity (Bray-Curtis) showing eDNA samples (n=59), with colors indicating sampling site by river kilometer (distance from confluence with the Yolo Bypass) and shapes indicating sampling season; see Table 1 for site information.

### Comparison with conventional sampling

Assemblages recovered from eDNA closely resembled conventional assemblages, though with some key differences. First, in paired sampling, eDNA always detected more species (Figures 4 and 5; Supplementary Figures S3 and S4). This pattern was statistically significant when eDNA detection was compared with concordant electrofishing samples (p < 0.01), with eDNA detecting an average of 4.6 more species per sampling event. eDNA detected an average of 5.3 more species per sampling event than concordant trap surveys (significance was not tested due to the small sample size). Only the higher effort multiple method survey detected a similar number of species compared with concurrent eDNA sampling, with 5 species unique to eDNA samples and 4 species (plus undetermined sunfishes) unique to the conventional aggregate method sample.

**Figure 4.**
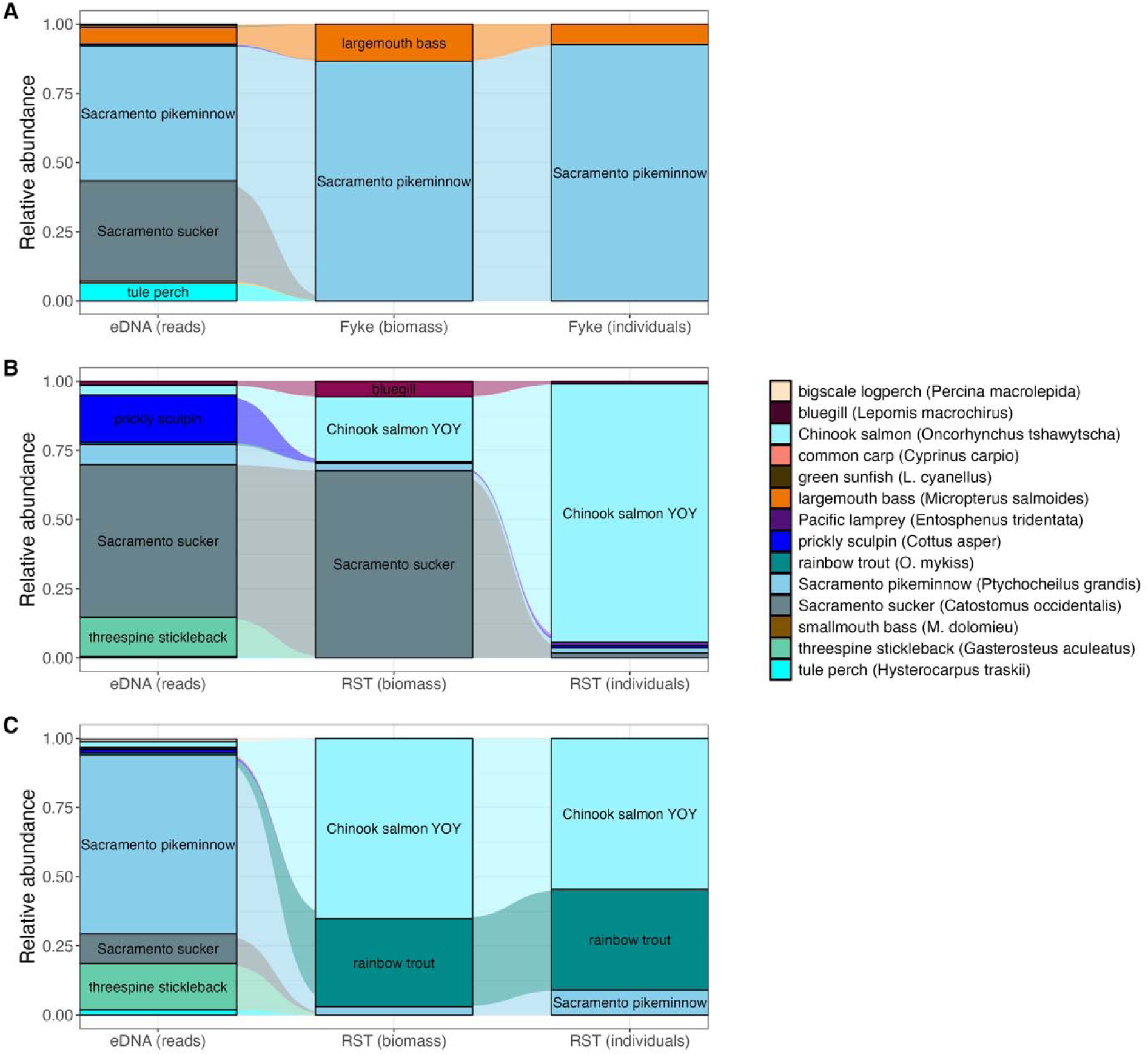
Alluvial plots comparing relative read abundance from eDNA with biomass and individual counts from the fyke trap at site RRAN in 2020 (A) and the rotary screw trap at site SW1D in April (B) and May (C) 2021; see Supplementary Figure S3 for corresponding Venn diagrams and Table 1 for site information.

**Figure 5.**
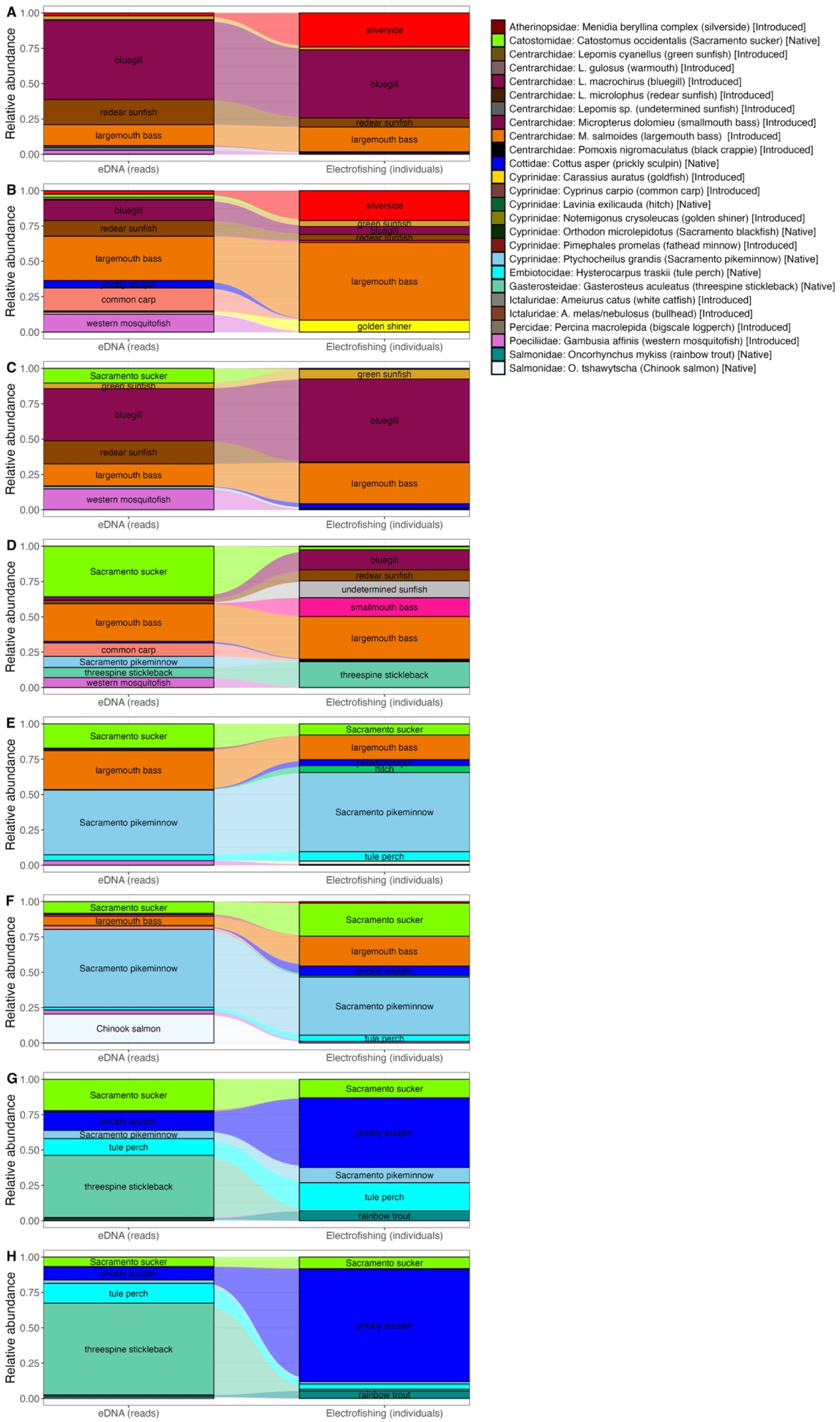
Alluvial plots comparing relative read abundance from eDNA with electrofishing counts: MACU in 2020 (A), MACU in 2021 (B), OLDR in 2020 (C), PEDR in 2020 (D), PEDR in 2021 (E), FRPG in 2021 (F; multi-method survey), I505 in 2021 (G), and WPCP in 2021 (H); see Supplementary Figure S4 for corresponding Venn diagrams and Table 1 for site information.

Species detected by eDNA but not the corresponding conventional surveys included non-native common carp, goldfish, and black crappie. Native Sacramento blackfish was not reported in any recent electrofishing surveys, but sequences identical to the Sacramento blackfish reference sequence were detected in eDNA samples not paired conventional surveys. Blackfish are common in the UC Davis arboretum, which flows over a small dam and empties into Putah Creek; thus, the species may be present in the watershed, but not the creek proper. Native Pacific lamprey and hitch were each reported once in trap and electrofishing samples, respectively, but not detected by concordant eDNA sampling. Non-native bigscale logperch detected by electrofishing 3 times but absent from concordant eDNA samples. However, bigscale logperch was detected once by eDNA and not by concordant electrofishing. One native species, Sacramento perch, and five non-native species reported from historical data (Jacinto et al., 2023) were not detected by eDNA or the concurrent conventional sampling in 2020-2021.

### Reordering and species-specific patterns

Modified rank-abundance curves (dominance-diversity Whittaker plots) show spatial reordering between native and non-native assemblages and indicate no apparent trends in evenness (Figure 6). Qualitative comparison of eDNA rank abundance curves with similar figures based on electrofishing data (Jacinto et al., 2023) show a longer “tail” on the eDNA curves, indicating that higher species richness in eDNA samples is primarily driven by increased sensitivity to rare taxa (Figure 6; Supplementary Figure S5). Consistent with long-term electrofishing data (Jacinto et al., 2023), more non-native species were detected at downstream sites and with higher relative read abundance, while native species dominated the assemblage of upstream, cold-water sites (Figure 6) and mean rank shift (MRS) values calculated at 1-year sampling intervals were higher at downstream sites (Supplementary Table S8; Supplementary Figure S6). Season-to-season MRS values were relatively higher compared with the year-to-year values at all sites other than the most upstream site. Species turnover was slightly higher at the most upstream site (Supplementary Table S8; Supplementary Figure S6). However, both MRS and turnover metrics should be interpreted cautiously, as they are generally more suitable for longer-term datasets.

**Figure 6.**
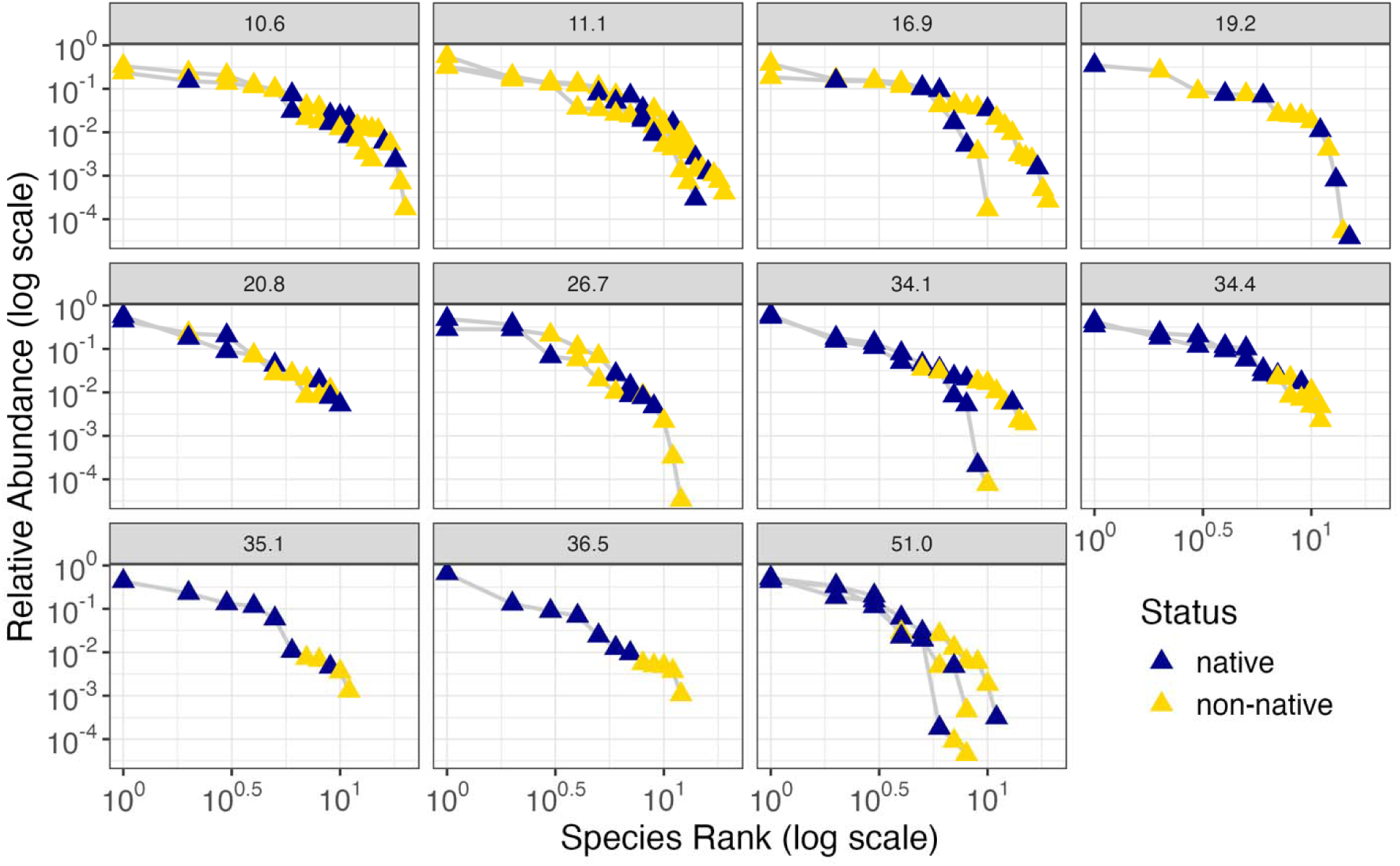
Rank-abundance curves (Whittaker plots) of native and non-native species identified by eDNA metabarcoding using relative sequence read abundance; plots are organized by river kilometer and each curve shows mean values from field replicates of a collecting event.

Finally, gradient forest analysis identified 13 species, six of which were native species, with significant associations (R² > 0.5) to spatial and temporal predictors (Figure 7). Among the species identified as significant, non-native bluegill and native three-spined stickleback showed the highest species importance scores. Five species were nonsignificant in the model. Eight species were not included in the model, likely because they were detected too infrequently to be reliably modeled. Among the three predictors, distance from confluence (km) had the greatest importance, followed by Julian date and year (Supplementary Figure S7).

**Figure 7.**
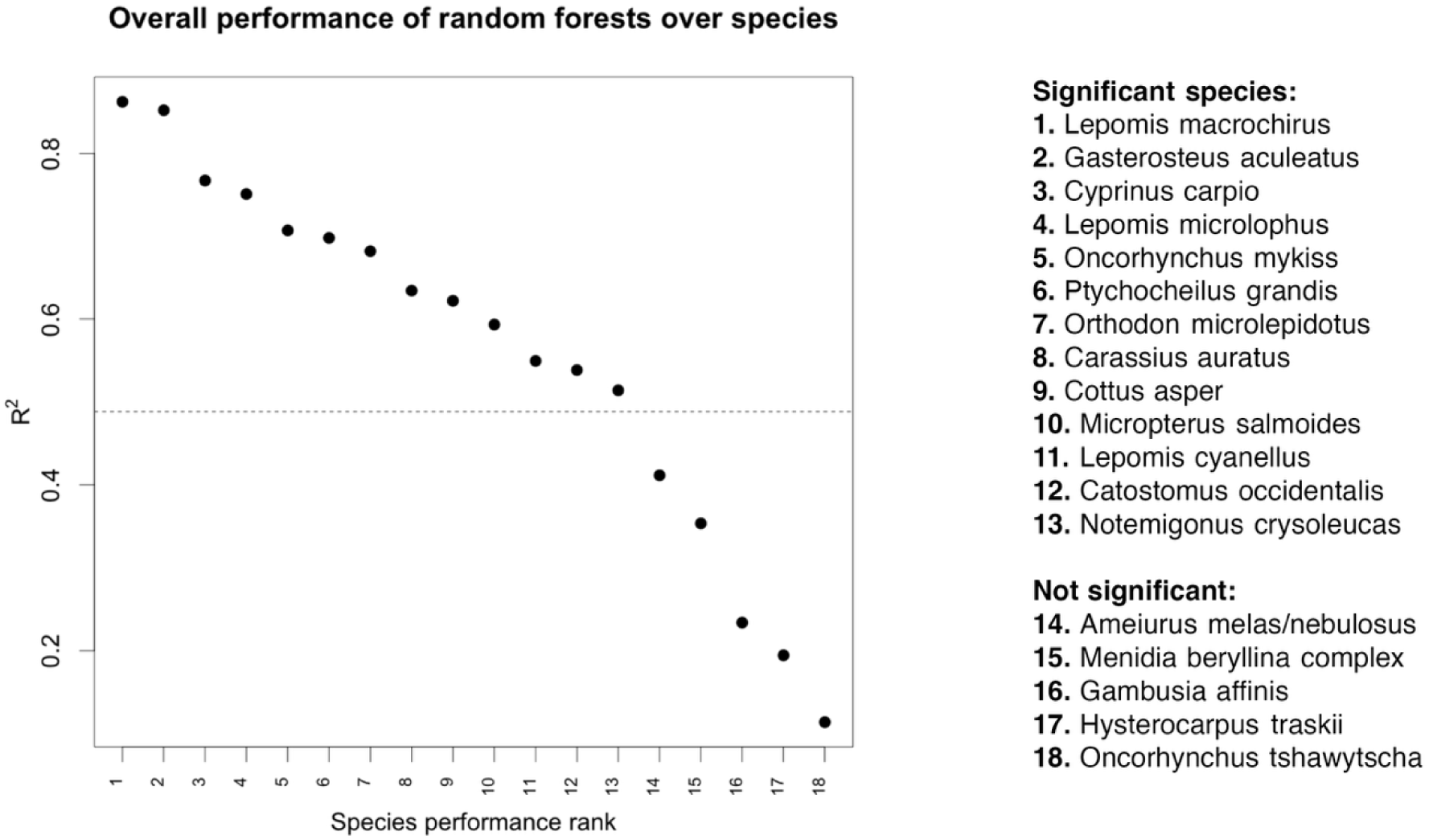
Gradient forest model showing the overall performance of individual species based on their response to predictor variables of river kilometer, sampling date, and year, with significance assessed using the broken stick method; predictor accuracy is provided in Supplementary Figure S6.

## Discussion

Dam infrastructure and hydrological alternations have significantly impacted freshwater fishes and continue to pose challenges worldwide. Despite growing investment in rehabilitation of streams and rivers, many efforts lack adequate monitoring to assess success (Bash & Ryan, 2002; Palmer et al., 2005). To address this gap, we examined whether eDNA metabarcoding can produce management-relevant measures of fish community diversity in a regulated stream. Overall, eDNA metabarcoding demonstrated repeatable detection of freshwater fishes and supported community level-analyses, highlighting its value as a complement to conventional fisheries monitoring.

Detections of both native and non-native species align with historical and concurrent fisheries monitoring using conventional sampling methods (Jacinto et al., 2023). Broad patterns of species composition reflected a transition from an upstream, coldwater assemblage to more diverse downstream assemblages typical of California’s Central Valley (May & Brown, 2002). Native species at upstream sites, including prickly sculpin, three-spined stickleback, and rainbow trout, were significant in the gradient forest model (Figure 7), indicating predictable relative abundance based on distance. eDNA captured the longitudinal pattern documented in electrofishing surveys (Jacinto et al. 2023) of relatively higher species richness in the lower reaches of the stream less influenced by cold water releases from the Putah Diversion Dam (PDD; Figures 3 and 4). In California, 39% of total species richness is attributed to non-native species (Mitchell & Knouft, 2009). Therefore, species richness is not necessarily a measure of ecological health, as disruptive non-natives dominate some of the most speciose reaches (e.g., common carp, bluegill, largemouth bass).

While ecological assemblages in natural streams are well-known to vary longitudinally because of landscape-level variation in stream size, riparian plant communities, and ecosystem ecology dynamics (Vannote et al., 1980; Gregory et al., 1991), observed variation in this fish assemblage is outside expected longitudinal changes for a stream of this size. Interestingly, central stream reaches below the PDD generally did not exhibit a gradual transition between communities, likely reflecting non-linear changes in local habitat conditions. Differences in species assemblages across a relatively limited spatial scale may reflect the influence of flow or local conditions like pools that provide refuge from predators which, in turn, influence freshwater fish community assembly (Moyle & Light, 1996).

Although the temporal scale of this study was limited, eDNA metabarcoding may be especially useful for understanding interannual and seasonal dynamics. PERMANOVA analysis using eDNA data identified community dissimilarity between spring and fall, a novel comparison for fish assemblages in this creek that was enabled by eDNA. Sampling date and year also contributed to gradient forest model accuracy, suggesting seasonal and interannual variation may also play meaningful roles in community structuring, like previous studies (Fitzgerald et al., 2017; Yeager et al., 2011). Mediterranean ecosystems are generally characterized by high seasonality which leads to substantial temporal variation in assemblages (Tonkin et al., 2017). However, annual electrofishing surveys can only be conducted in low flow while traps require high flow, making direct comparisons of high (spring) and low (fall) flow conditions biased by gear. Future monitoring could leverage the scalability of eDNA by implementing high frequency sampling, particularly when conventional sampling is constrained by flow conditions, cost, or other logistical challenges.

Because measures of fish community structure are strongly influenced by sampling effort (Angermeier & Smogor, 1995), the higher detection sensitivity of eDNA metabarcoding can provide new insight on community assembly dynamics (Seymour et al., 2021; this study). With just two years of eDNA data, we were able examine community dynamics such as rank abundance in a manner similar to those derived from 25 years of electrofishing surveys (Jacinto et al., 2023). The higher sensitivity of eDNA also facilitates the inclusion of relatively rarer species in analyses such as gradient forest models, which excludes species with insufficient data. Seasonal differences between spring and fall fish communities were also evident, a result not detectable with conventional sampling because different gears are deployed according to seasonal flow conditions.

In addition to corroborating previously documented patterns of native species recovery and facilitating community-level analyses, eDNA was particularly effective at detecting two species that were underrepresented in concurrent conventional surveys, likely due to small size or low abundance. eDNA was more effective for detection of three-spined stickleback, a small fish for which electrofishing efficiency is low (Penczak & Głowacki, 2008). eDNA also detected common carp when they were not observed in concurrent electrofishing surveys, indicating that barge electrofishing may be ineffective for detecting carp in low-density populations. Effective electrofishing detection of invasive freshwater carps can require refined techniques (Bouska et al., 2017) and increased target power (Young et al., 2023).

Discrepancies between eDNA and conventional survey data may result from detection biases and eDNA transport. For example, bigscale logperch is a small, benthic species that was underrepresented in eDNA data and may be hard to detect in water collected near the surface. Pacific lamprey, which live in riparian systems as benthic juveniles (ammocoetes), are also poorly amplified by MiFish primers (Miya et al., 2020). Detection rates of these species could be improved by increasing the number of field replicates, increasing water sampling depth, or using multiple eDNA metabarcoding markers (Li et al., 2018). Detections of kokanee, threadfin shad, and Chinook salmon eDNA at Stebbins Cold Canyon, the most upstream site, is consistent with limited downstream transport of eDNA (Pont et al., 2018) from Lake Berryessa.

Taxonomic challenges may also complicate interpretation of eDNA metabarcoding results. While the MiFish marker provided species-level resolution for most taxa in this study, short barcodes do not always resolve closely related species (Yates et al., 2025). Although salmonids and non-native minnow species pairs had the lowest genetic distances (<3%; Supplementary Table S6), the ASVs of these seven species pairs were consistently distinguishable. This suggests that clustering sequences into operational taxonomic units (OTUs) based on similarity thresholds of 97–99% would have obscured taxonomically informative sequence variation. For bullheads (Dumke et al., 2020), minnows (April et al. 2011), and sculpins (Moyle & Campbell, 2022) in the Putah Creek assemblage, evolutionary complexity can contribute to difficulty identifying these species both morphologically and using eDNA.

Additionally, hybridization has been documented within bullheads (Walter et al., 2014), sunfishes (Kiernan et al. 2012), and some native minnows (Baumsteiger & Moyle, 2019; Moyle & Massingill, 1981). Because 12S rRNA is a mitochondrial gene, any hybrids detected using eDNA would have to be assigned according to their maternal lineage; the paternal lineage cannot be determined from this marker. Conversely, eDNA metabarcoding can also reveal unrecognized genetic variation; the detection of a second prickly sculpin haplotype was unexpected given *Cottus* spp. are typically difficult to differentiate using short barcodes (Yates et al., 2025). While additional work is needed to understand this result, cryptic genetic diversity would be consistent with the complex evolutionary histories of California freshwater sculpins (Moyle & Campbell, 2022).

Although conservation efforts often focus on species richness, other measures such as relative abundance (Wilsey et al., 2005) and reordering (Hillebrand et al., 2018) may be more meaningful indicators of biodiversity change, especially in response to management action or human disturbance (Collins et al., 2008). Our results are consistent with a weakly quantitative relationship between relative sequence abundance and fish biomass (Lamb et al., 2019) that best reflects the relative abundance of dominant species (Skelton et al., 2023). Although methodological biases do have the potential to obscure real differences in relative abundances of species (Kelly et al., 2019), considering eDNA metabarcoding as semi-quantitative (rather than presence/absence) enhances the resolution of community-level differences (Guri et al., 2023).

Reordering is recognized as an important and under-studied spatiotemporal dynamic that provides ecological insight in both terrestrial (e.g., Jones et al., 2017; Avolio et al., 2019) and aquatic (e.g., Jacinto et al., 2023) systems and can be a central driver of long-term community changes (Jones et al., 2017). Reordering (including but not limited to species turnover) may be an important dynamic in Putah Creek, and studies expanding mean rank shift (MRS) comparisons across seasons to more sites could help inform management of environmental flow. MRS is an indicator of assemblage stability, and the exploratory values calculated here showed higher season to season MRS compared with interannual values (Figure 7); this dynamic cannot not be captured when surveys are conducted once a year. Spatiotemporal patterns in species turnover were less evident; relatively higher upstream values may reflect typical seasonal variation characteristic of native assemblages or seasonal flow from Lake Berryessa but should be interpreted cautiously.

The concordance between eDNA and conventional survey data (Jacinto et al., 2023) in community stability metrics like MRS provides a degree of confidence in the utility of this application of eDNA data and suggest potential future uses for management. Metrics like MRS align with management frameworks like Putah Creek’s, based on California Fish and Game Code 5937, which stipulates that fish populations in streams below dams be kept in ‘good condition.’ Future restoration work in Putah Creek will likely focus on enhancing the habitat and ecological stability of the lower portions of the watershed. While eDNA values should be interpreted with caution due to the uncertainty in the quantitative ability of metabarcoding and the assumption that meaningful ecological change takes place over longer time intervals, eDNA has unique potential to provide insight where conventional sampling cannot (i.e., due to seasonal flows requiring different sampling techniques). This study demonstrates how eDNA detection data, now increasingly available, might be used to answer more basic and applied ecological questions, including those related to assembly dynamics.

Freshwater fishes are indicators of stream health and are ecologically and culturally valuable (Pander & Geist, 2013; Reid et al., 2021; Rypel et al., 2021); conservation and management require effective monitoring of community responses. The patterns observed in this study align with our understanding of California’s natural flow regime that overall favors native species (Moyle 2014) and discourages non-native species (Moyle & Mount, 2007) and add to a growing body of work demonstrating how environmental flows promote recovery of native fish (Baruch et al., 2024; Hitt et al., 2025; Jacinto et al., 2023). While conventional surveys also offer valuable insight that eDNA cannot (e.g., fish size, condition), repeated sampling is not always feasible. eDNA metabarcoding can be more cost-effective than conventional methods for multi-species detection (Lafferty, 2024), making it particularly suitable for long-term monitoring, surveys across heterogeneous conditions, and logistically challenging habitats. The use of eDNA to improve our understanding of fish community responses to stream rehabilitation could help guide recovery efforts of native species (Bennett et al., 2016; Pander & Geist, 2013; Roni et al., 2018). For example, eDNA is currently being implemented to track community responses to Klamath River Renewal Project, the largest dam removal and river restoration in California and the United States. This project has established molecular reference specimens and baseline biodiversity assessments using eDNA metabarcoding (Keel et al., 2025).

## Conclusion

Our findings demonstrate the applicability of eDNA metabarcoding approaches for stream ecology and restoration science. In this study, eDNA provided management relevant insight into fish community changes across space and time, consistently detected more species than conventional methods, and enabled repeated sampling across changing river conditions. As methods and resources (e.g., regional reference sequence databases) are developed and optimized, eDNA can be a cost-effective and efficient method for fish community assessment.

Beyond species detection, these results support the use of eDNA metabarcoding to quantify community-level ecological dynamics including assembly, stability, and reordering. These community dynamics can be incorporated into whole-ecosystem approaches to ecosystem management, which may be more effective than single-species approaches (Moyle et al., 2017; Mount et al., 2019; Obester et al., 2022). A more nuanced understanding of the assembly dynamics of riverine fish assemblages could help facilitate management strategies that help preserve imperiled species and freshwater biodiversity. Insights into community dynamics gleaned from eDNA metabarcoding can inform conservation and management of declining and fragile freshwater fauna at all scales.

## Acknowledgments

We thank Solano County Water Agency (Contract 03-00206VR) for primary support of this project. AEH gratefully acknowledges additional support from the Diablo Valley Flyfishing Club, California Fly Fishers Unlimited, Marin Rod and Gun Club, and Fly Fishers of Davis.

Sequencing was carried out at the DNA Technologies and Expression Analysis Core at the UC Davis Genome Center, supported by NIH Shared Instrumentation Grant 1S10OD010786-01. We also thank Tim Salamunovich and TRPA Fish Biologists for the electrofishing survey data, local landowners who provided access to Putah Creek for sampling, Khyana Yearwood and Shayan Kaveh for assistance with field sampling and laboratory work, and Ted Grosholz for feedback that helped improve this manuscript. NAF and ALR were supported by the University of California, Davis Agricultural Experiment Station (CA-D-WFB2098-H, CA-D-WFB2467-H, respectively), and ALR was also supported by the Peter B. Moyle and California Trout Endowment for Coldwater Fish Conservation.

## References

Altschul, S. F., W. Gish, W. Miller, E. W. Myers, and D. J. Lipman. 1990. “Basic Local Alignment Search Tool.” Journal of Molecular Biology 215(3): 403–10. 10.1016/S0022-2836(05)80360-2.

Angermeier, P. L., and R. A. Smogor. 1995. “Estimating Number of Species and Relative Abundances in Stream-Fish Communities: Effects of Sampling Effort and Discontinuous Spatial Distributions.” Canadian Journal of Fisheries and Aquatic Sciences 52(5): 936–949. 10.1139/f95-093.

Arthington, A. H., N. K. Dulvy, W. Gladstone, and I. J. Winfield. 2016. “Fish Conservation in Freshwater and Marine Realms: Status, Threats and Management.” Aquatic Conservation: Marine and Freshwater Ecosystems 26(5): 838–857. 10.1002/aqc.2712.

Avolio, M. L., I. T. Carroll, S. L. Collins, G. R. Houseman, L. M. Hallett, F. Isbell, S. E. Koerner, K. J. Komatsu, M. D. Smith, and K. R. Wilcox. 2019. “A Comprehensive Approach to Analyzing Community Dynamics Using Rank Abundance Curves.” Ecosphere 10(10): e02881. 10.1002/ecs2.2881.

Baruch, E. M., S. M. Yarnell, T. E. Grantham, J. R. Ayers, A. L. Rypel, and R. A. Lusardi. 2024. “Mimicking Functional Elements of the Natural Flow Regime Promotes Native Fish Recovery in a Regulated River.” Ecological Applications 34(6): e3013. 10.1002/eap.3013.

Bash, J. S., and C. M. Ryan. 2002. “Stream Restoration and Enhancement Projects: Is Anyone Monitoring?” Environmental Management 29(6): 877–885. 10.1007/s00267-001-0066-3.

Baumsteiger, J., and P. B. Moyle. 2019. “A Reappraisal of the California Roach/Hitch (Cypriniformes, Cyprinidae, Hesperoleucus/Lavinia) Species Complex.” Zootaxa 4543(2): 221–240. 10.11646/zootaxa.4543.2.3.

Benke, A. C. 1990. “A Perspective on America’s Vanishing Streams.” Journal of the North American Benthological Society 9(1): 77–88. 10.2307/1467936.

Bennett, S., G. Pess, N. Bouwes, P. Roni, R. E. Bilby, S. Gallagher, J. Ruzycki, T. Buehrens, et al. 2016. “Progress and Challenges of Testing the Effectiveness of Stream Restoration in the Pacific Northwest Using Intensively Monitored Watersheds.” Fisheries 41(2): 92–103. 10.1080/03632415.2015.1127805.

Betancur-R, R., E. O. Wiley, G. Arratia, A. Acero, N. Bailly, M. Miya, G. Lecointre, and G. Ortí. 2017. “Phylogenetic Classification of Bony Fishes.” BMC Evolutionary Biology 17: 162. 10.1186/s12862-017-0958-3.

Börk, K. S., J. F. Krovoza, J. V. Katz, and B. Peter. 2012. “The Rebirth of California Fish and Game Code Section 5937: Water for Fish.” University of California Davis Law Review 45: 809–913.

Bouska, W. W., D. C. Glover, K. L. Bouska, and J. E. Garvey. 2017. “A Refined Electrofishing Technique for Collecting Silver Carp: Implications for Management.” North American Journal of Fisheries Management 37(1): 101–107. 10.1080/02755947.2016.1240122.

Brown, L. R., and M. L. Bauer. 2010. “Effects of Hydrologic Infrastructure on Flow Regimes of California’s Central Valley Rivers: Implications for Fish Populations.” River Research and Applications 26(6): 751–765. 10.1002/rra.1293.

Brunson, J. C., and Q. D. Read. 2023. “ggalluvial: Alluvial Plots in ‘ggplot2.’” R package version 0.12.5. https://corybrunson.github.io/ggalluvial/.

Bunn, S. E., and A. H. Arthington. 2002. “Basic Principles and Ecological Consequences of Altered Flow Regimes for Aquatic Biodiversity.” Environmental Management 30(4): 492–507. 10.1007/s00267-002-2737-0.

Callahan, B. J., P. J. McMurdie, M. J. Rosen, A. W. Han, A. J. A. Johnson, and S. P. Holmes.2016. “DADA2: High-Resolution Sample Inference from Illumina Amplicon Data.” Nature Methods 13(7): 581–583. 10.1038/nmeth.3869.

Collins, S. L., K. N. Suding, E. E. Cleland, M. Batty, S. C. Pennings, K. L. Gross, J. B. Grace, L. Gough, J. E. Fargione, and C. M. Clark. 2008. “Rank Clocks and Plant Community Dynamics.” Ecology 89(12): 3534–3541. 10.1890/07-1646.1.

Diver, S., J. R. Oberholzer Dent, D. Sarna-Wojcicki, R. Reed, and C. Dill-De Sa. 2024. “Recasting Klamath Dam Removal as Eco-Cultural Revitalization and Restorative Justice through Karuk Tribal Leadership.” Water 16(16): 2295. 10.3390/w16162295.

Dumke, J. D., G. M. Chorak, C. R. Ruetz, R. A. Thum, and J. N. Wesolek. 2020. “Identification of Black Bullhead (*Ameiurus melas*) and Brown Bullhead (*A. nebulosus*) from the Western Great Lakes: Recommendations for Small Individuals.” The American Midland Naturalist 183(1): 90–104. 10.1637/19-041.

Dynesius, M., and C. Nilsson. 1994. “Fragmentation and Flow Regulation of River Systems in the Northern Third of the World.” Science 266(5186): 753–762. 10.1126/science.266.5186.753.

Ellis, N., S. J. Smith, and C. R. Pitcher. 2012. “Gradient Forests: Calculating Importance Gradients on Physical Predictors.” Ecology 93(1): 156–168. 10.1890/11-0252.1.

Evans, N. T., P. D. Shirey, J. G. Wieringa, A. R. Mahon, and G. A. Lamberti. 2017. “Comparative Cost and Effort of Fish Distribution Detection via Environmental DNA Analysis and Electrofishing.” Fisheries 42(2): 90–99. 10.1080/03632415.2017.1276329.

Feng, X., B. Li, Y. Chen, R. Zhu, Y. Jia, and X. Sui. 2023. “Species-level Monitoring of Rare and Invasive Fishes using eDNA Metabarcoding in the Middle and Upper Yarlung Zangbo River, Tibet.” Water Biology and Security, 2(1), 100089. 10.1016/j.watbs.2022.100089.

Fitzgerald, D. B., K. O. Winemiller, M. H. Sabaj Pérez, and L. M. Sousa. 2017. “Seasonal Changes in the Assembly Mechanisms Structuring Tropical Fish Communities.” Ecology 98(1): 21–31. 10.1002/ecy.1612.

Fluker, B. L., F. Pezold, and R. L. Minton. 2011. “Molecular and Morphological Divergence in the Inland Silverside (Menidia beryllina) along a Freshwater-Estuarine Interface.” Environmental Biology of Fishes 91(3): 311–325. 10.1007/s10641-011-9786-2.

Folke, C. 2003. “Freshwater for Resilience: A Shift in Thinking.” Philosophical Transactions of the Royal Society of London. Series B: Biological Sciences 358(1440): 2027–2036. 10.1098/rstb.2003.1385.

Frontier, S. 1976. “Étude de la Décroissance des Valeurs Propres dans une Analyse en Composantes Principales: Comparaison avec le Modèle du Bâton Brisé.” Journal of Experimental Marine Biology and Ecology 25(1): 67–75. 10.1016/0022-0981(76)90076-9.

Gleick, P. H., and H. Cooley. 2021. “Freshwater Scarcity.” Annual Review of Environment and Resources 46(1): 319–348. 10.1146/annurev-environ-012220-101319.

Goldberg, C. S., C. R. Turner, K. Deiner, K. E. Klymus, P. F. Thomsen, M. A. Murphy, S. F. Spear, et al. 2016. “Critical Considerations for the Application of Environmental DNA Methods to Detect Aquatic Species.” Methods in Ecology and Evolution 7(11): 1299–1307. 10.1111/2041-210X.12595.

Gregory, S. V., F. J. Swanson, W. A. McKee, and K. W. Cummins. 1991. “An Ecosystem Perspective of Riparian Zones.” BioScience 41(8): 540–551. 10.2307/1311607.

Gudde, E. G. 2004. California Place Names: The Origin and Etymology of Current Geographical Names. 4th ed., revised and enlarged by W. Bright. University of California Press.

Guri, G., I. Westgaard, N. Yoccoz, O. S. Wangensteen, K. Præbel, J. Louise, R. Ryan, A. P. K., O. Shelton, T. Hanebrekke, and T. Johansen. 2023. “Maximizing Sampling Efficiency to Detect Differences in Fish Community Composition Using Environmental DNA Metabarcoding in Subarctic Fjords.” Environmental DNA 6(1): e409. 10.1002/edn3.409.

Hallett, L. M., S. K. Jones, A. A. M. MacDonald, M. B. Jones, D. F. B. Flynn, J. Ripplinger, P. Slaughter, C. Gries, and S. L. Collins. 2016. “codyn: An R Package of Community Dynamics Metrics.” Methods in Ecology and Evolution 7(10): 1146–1151. 10.1111/2041-210X.12569.

Hallett, L., M. Avolio, I. Carroll, S. Jones, A. MacDonald, D. Flynn, P. Slaughter, J. Ripplinger, S. Collins, C. Gries, and M. Jones. 2020. *codyn: Community Dynamics Metrics*. R package version 2.0.5. 10.5063/F1N877Z6.

Hänfling, B., L. Lawson Handley, D. S. Read, C. Hahn, J. Li, P. Nichols, R. C. Blackman, A. Oliver, and I. J. Winfield. 2016. “Environmental DNA Metabarcoding of Lake Fish Communities Reflects Long-Term Data from Established Survey Methods.” Molecular Ecology 25(13): 3101–3119. 10.1111/mec.13660.

Harrison, I., R. Abell, W. Darwall, M. L. Thieme, D. Tickner, and I. Timboe. 2018. “The Freshwater Biodiversity Crisis.” Science 362(6421): 1369. 10.1126/science.aav9242.

Heino, J., R. Virkkala, and H. Toivonen. 2009. “Climate Change and Freshwater Biodiversity: Detected Patterns, Future Trends and Adaptations in Northern Regions.” Biological Reviews 84(1): 39–54. 10.1111/j.1469-185X.2008.00060.x.

Hillebrand, H., B. Blasius, E. T. Borer, J. M. Chase, J. A. Downing, B. K. Eriksson, C. T. Filstrup, et al. 2018. “Biodiversity Change Is Uncoupled from Species Richness Trends: Consequences for Conservation and Monitoring.” Journal of Applied Ecology 55(1): 169–184. 10.1111/1365-2664.12959.

Hitt, L. G., M. Willmes, G. Whitman, M. C. Miner, C. Jeffres, R. C. Johnson, … A. L. Rypel. 2025. “Early Evidence for Establishment of a Chinook Salmon Population in a Restored Watershed.” Ecosphere 16(3): e70207. 10.1002/ecs2.70207.

Holmes, A. E., M. R. Baerwald, J. Rodzen, B. M. Schreier, B. Mahardja, and A. J. Finger. 2024. “Evaluating Environmental DNA Detection of a Rare Fish in Turbid Water Using Field and Experimental Approaches.” PeerJ 12: e16453. 10.7717/peerj.16453.

Illumina. 2013. 16S Metagenomic Sequencing Library Preparation (Document No. 15044223 B). Illumina, Inc. https://support.illumina.com/downloads/16s_metagenomic_sequencing_library_preparation.html.

Jacinto, E., N. A. Fangue, D. E. Cocherell, J. D. Kiernan, P. B. Moyle, and A. L. Rypel. 2023. “Increasing Stability of a Native Freshwater Fish Assemblage Following Flow Rehabilitation.” Ecological Applications 33(5): e2868. 10.1002/eap.2868.

Jones, S. K., J. Ripplinger, and S. L. Collins. 2017. “Species Reordering, Not Changes in Richness, Drives Long-Term Dynamics in Grassland Communities.” Ecology Letters 20(12): 1556–1565. 10.1111/ele.12864.

Keel, D. J., K. Karpenko, S. M. Blankenship, G. Schumer, O. O’Rourke, C. O. Ostberg, D. A. Chase, and J. J. Duda. 2025. “A Molecular Specimen Bank for Contemporary and Future Study Captures Landscape-Scale Biodiversity Baselines Before Klamath River Dam Removal.” Scientific Reports 15(1): 20679. 10.1038/s41598-025-07042-1

Kelly, R. P., A. O. Shelton, and R. Gallego. 2019. “Understanding PCR Processes to Draw Meaningful Conclusions from Environmental DNA Studies.” Scientific Reports 9(1): 1–14. 10.1038/s41598-019-48546-x.

Kiernan, J. D., P. B. Moyle, and P. K. Crain. 2012. “Restoring Native Fish Assemblages to a Regulated California Stream Using the Natural Flow Regime Concept.” Ecological Applications 22(5): 1472–1482. 10.1890/11-0480.1.

Konrad, C. P., J. D. Olden, D. A. Lytle, T. S. Melis, J. C. Schmidt, E. N. Bray, M. C. Freeman, et al. 2011. “Large-Scale Flow Experiments for Managing River Systems.” BioScience 61(12): 948–959. 10.1525/bio.2011.61.12.5.

Lafferty, K. 2024. “Metabarcoding Is (Usually) More Cost Effective than Seining or qPCR for Detecting Tidewater Gobies and Other Estuarine Fishes.” PeerJ 12: e16847. 10.7717/peerj.16847.

Lamb, P. D., E. Hunter, J. K. Pinnegar, S. Creer, R. G. Davies, and M. I. Taylor. 2019. “How Quantitative Is Metabarcoding: A Meta-Analytical Approach.” Molecular Ecology 28: 420–430. 10.1111/mec.14920.

Lejon, A. G. C., B. Malm Renöfält, and C. Nilsson. 2009. “Conflicts Associated with Dam Removal in Sweden.” Ecology and Society 14(2): 4. 10.5751/ES-02931-140204.

Li, Y., N. T. Evans, M. A. Renshaw, C. L. Jerde, B. P. Olds, A. J. Shogren, K. Deiner, D. M. Lodge, G. A. Lamberti, and M. E. Pfrender. 2018. “Estimating Fish Alpha- and Beta-Diversity along a Small Stream with Environmental DNA Metabarcoding.” Metabarcoding and Metagenomics 2: e24262. 10.3897/mbmg.2.24262.

Lund, J. 2016. “California’s Agricultural and Urban Water Supply Reliability and the Sacramento–San Joaquin Delta.” San Francisco Estuary and Watershed Science 14(3): 6. 10.15447/sfews.2016v14iss3art6.

Lynch, A. J., S. J. Cooke, A. H. Arthington, C. Baigun, L. Bossenbroek, C. Dickens, I. Harrison, et al. 2023. “People Need Freshwater Biodiversity.” WIREs Water 10(3): e1633. 10.1002/wat2.1633.

Lytle, D. A., and N. L. Poff. 2004. “Adaptation to Natural Flow Regimes.” Trends in Ecology & Evolution 19(2): 94–100. 10.1016/j.tree.2003.10.002.

Martin, M. 2011. “Cutadapt Removes Adapter Sequences from High-Throughput Sequencing Reads.” EMBnet.journal 17(1): 10. 10.14806/ej.17.1.200.

May, J. T., and L. R. Brown. 2002. “Fish Communities of the Sacramento River Basin: Implications for Conservation of Native Fishes in the Central Valley, California.” Environmental Biology of Fishes 63(4): 373–388. 10.1023/A:1020877717701.

McElroy, M. E., T. L. Dressler, G. C. Titcomb, E. A. Wilson, K. Deiner, T. L. Dudley, E. J. Eliason, et al. 2020. “Calibrating Environmental DNA Metabarcoding to Conventional Surveys for Measuring Fish Species Richness.” Frontiers in Ecology and Evolution 8: 276. 10.3389/fevo.2020.00276.

McMurdie, P. J., and S. Holmes. 2013. “phyloseq: An R Package for Reproducible Interactive Analysis and Graphics of Microbiome Census Data.” PLoS ONE 8(4): e61217. 10.1371/journal.pone.0061217.

Miner, M. 2022. “Migratory Phenology and Spatial Distributions of a Recovering Chinook Salmon Run in a Flow Regulated Creek, Considerations for Management.” Master’s thesis, University of California, Davis, California, USA.

Mitchell, A. L., and J. H. Knouft. 2009. “Non-Native Fishes and Native Species Diversity in Freshwater Fish Assemblages across the United States.” Biological Invasions 11(6): 1441– 1450. 10.1007/s10530-008-9352-9.

Miya, M., Y. Sato, T. Fukunaga, T. Sado, J. Y. Poulsen, K. Sato, T. Minamoto, et al. 2015. “MiFish, a Set of Universal PCR Primers for Metabarcoding Environmental DNA from Fishes: Detection of More than 230 Subtropical Marine Species.” Royal Society Open Science 2(7): 150088. 10.1098/rsos.150088.

Miya, M., R. O. Gotoh, and T. Sado. 2020. “MiFish Metabarcoding: A High Throughput Approach for Simultaneous Detection of Multiple Fish Species from Environmental DNA and Other Samples.” Fisheries Science 86: 939–970. 10.1007/s12562-020-01461-x.

Mount, J., B. Gray, K. Bork, J. Cloern, F. Davis, T. Grantham, L. Grenier, J. L. Harder, Y. Kuwayama, P. Moyle, and M. Schwartz. 2019. A Path Forward for California’s Freshwater Ecosystems. Public Policy Institute of California, San Francisco, California, USA. https://www.ppic.org/publication/a-path-forward-for-californias-freshwater-ecosystems/.

Moyle, P. B. 2014. “Novel Aquatic Ecosystems: The New Reality for Streams in California and Other Mediterranean Climate Regions.” River Research and Applications 30: 1335–1344. 10.1002/rra.2709.

Moyle, P. B., and M. A. Campbell. 2022. “Cryptic Species of Freshwater Sculpin (Cottidae: *Cottus*) in California, USA.” Zootaxa 5154(5): 501–527. 10.11646/zootaxa.5154.5.1.

Moyle, P. B., J. V. Katz, and R. M. Quiñones. 2011. “Rapid Decline of California’s Native Inland Fishes: A Status Assessment.” Biological Conservation 144(10): 2414–2423. 10.1016/j.biocon.2011.06.002.

Moyle, P. B., and T. Light. 1996. “Fish Invasions in California: Do Abiotic Factors Determine Success?” Ecology 77(6): 1666–1670. 10.2307/2265770.

Moyle, P. B., M. P. Marchetti, J. Baldrige, and T. L. Taylor. 1998. “Fish Health and Diversity: Justifying Flows for a California Stream.” Fisheries 23(7): 6–15. 10.1577/1548-8446(1998)023<0006:fhadjf>2.0.co;2.

Moyle, P. B., and M. Massingill. 1981. “Hybridization Between Hitch, *Lavinia exilicauda*, and Sacramento Blackfish, *Orthodon microlepidotus*, in San Luis Reservoir, California.” California Fish and Game 67: 196–198.

Moyle, P. B., and J. F. Mount. 2007. “Homogenous Rivers, Homogenous Faunas.” Proceedings of the National Academy of Sciences 104(14): 5711–5712. 10.1073/pnas.070145710.

Nagarajan, R., A. Holmes, and A. Schreier. 2023. Reference sequence database for eDNA metabarcoding of San Francisco estuary fishes and invertebrates [Dataset]. Dryad. 10.5061/dryad.0p2ngf25z.

Null S. E., J. Medellín-Azuara, A. Escriva-Bou, M. Lent, and J. R. Lund. 2014. “Optimizing the Dammed: Water Supply Losses and Fish Habitat Gains from Dam Removal in California.” Journal of Environmental Management 136: 121–131. 10.1016/j.jenvman.2014.01.024.

Obester, A. N., R. A. Lusardi, N. R. Santos, R. A. Peek, and S. M. Yarnell. 2022. “The Use of Umbrella Fish Species to Provide a More Comprehensive Approach for Freshwater Conservation Management.” Aquatic Conservation: Marine and Freshwater Ecosystems 32: 112–128. 10.1002/aqc.3746.

Oksanen, J., F. G. Blanchet, M. Friendly, R. Kindt, P. Legendre, D. McGlinn, P. R. Minchin, et al. 2022. “Vegan: Community Ecology Package.” R Package Version 2.6–4. https://CRAN.R-project.org/package=vegan.

Page, L. M., H. Espinosa-Pérez, L. T. Findley, C. R. Gilbert, R. N. Lea, N. E. Mandrak, M. d. M. Espinosa-García, and C. W. Becker. 2023. Common and Scientific Names of Fishes from the United States, Canada, and Mexico, 8^th^ ed. Bethesda, MD: American Fisheries Society.

Palmer, M. A., E. S. Bernhardt, J. D. Allan, P. S. Lake, G. Alexander, S. Brooks, J. Carr, et al. 2005. “Standards for Ecologically Successful River Restoration.” Journal of Applied Ecology 42: 208–217. 10.1111/j.1365-2664.2005.01004.x.

Pander, J., and J. Geist. 2013. “Ecological Indicators for Stream Restoration Success.” Ecological Indicators 30: 106–118. 10.1016/j.ecolind.2013.01.039.

Paradis E., and Schliep K. 2019. “ape 5.0: an environment for modern phylogenetics and evolutionary analyses in R.” Bioinformatics 35: 526–528. 10.1093/bioinformatics/bty633.

Parisek, C. A., F. A. De Castro, J. D. Colby, G. R. Leidy, S. Sadro, and A. L. Rypel. 2024. “Reservoir Ecosystems Support Large Pools of Fish Biomass.” Scientific Reports 14: 9428. 10.1038/s41598-024-59730-z.

Penczak, T., and Ł. Głowacki. 2008. “Evaluation of Electrofishing E ciency in a Stream Under Natural and Regulated Conditions.” Aquatic Living Resources 21: 329–337. 10.1051/alr:2008044.

Pinter, N., J. Lund, and P. Moyle. 2019. “The California Water Model: Resilience through Failure.” Hydrological Processes 33: 1775–1779. 10.1002/hyp.13447.

Poff, N. L., J. D. Allan, M. B. Bain, J. R. Karr, K. L. Prestegaard, B. D. Richter, R. E. Sparks, and J. C. Stromberg. 1997. “The Natural Flow Regime: A Paradigm for River Conservation and Restoration.” BioScience 47: 769–784. 10.2307/1313099.

Pokhrel, Y., M. Burbano, J. Roush, H. Kang, V. Sridhar, and D. Hyndman. 2018. “A Review of the Integrated Effects of Changing Climate, Land Use, and Dams on Mekong River Hydrology.” Water 10: 266. 10.3390/w10030266.

Pont, D., M. Rocle, A. Valentini, R. Civade, P. Jean, A. Maire, N. Roset, M. Schabuss, H. Zornig, and T. Dejean. 2018. “Environmental DNA Reveals Quantitative Patterns of Fish Biodiversity in Large Rivers Despite Its Downstream Transportation.” Scientific Reports 8: 10361. 10.1038/s41598-018-28424-8.

Price, A. L., and J. T. Peterson. 2010. “Estimation and Modeling of Electrofishing Capture Efficiency for Fishes in Wadeable Warmwater Streams.” North American Journal of Fisheries Management 30: 481–498. 10.1577/m09-122.1.

Reid, A. J., L. E. Eckert, J. Lane, N. Young, S. G. Hinch, C. T. Darimont, S. J. Cooke, N. C. Ban, and A. Marshall. 2021. “Two Eyed Seeing: An Indigenous Framework to Transform Fisheries Research and Management.” Fish and Fisheries 22: 243–261. 10.1111/faf.12516.

Richter, B. D., R. Mathews, D. L. Harrison, and R. Wigington. 2003. “Ecologically Sustainable Water Management: Managing River Flows for Ecological Integrity.” Ecological Applications 13: 206–224. 10.1890/1051-0761(2003)013[0206:ESWMMR]2.0.CO;2.

da Rocha, M., R. L. de Assis, M. T. F. Piedade, Y. O. Feitosa, J. E. Householder, G. D. S. Lobo, L. O. Demarchi, et al. 2019. “Thirty Years after Balbina Dam: Diversity and Floristic Composition of the Downstream Floodplain Forest, Central Amazon, Brazil.” Ecohydrology 12: e2144. 10.1002/eco.2144.

Roni, P., U. Åberg, and C. Weber. 2018. “A Review of Approaches for Monitoring the Effectiveness of Regional River Habitat Restoration Programs.” North American Journal of Fisheries Management 38: 1170–1186. 10.1002/nafm.10222.

Rosenzweig, M. L. 2003. “Reconciliation Ecology and the Future of Species Diversity.” Oryx 37: 194–205. 10.1017/S0030605303000371.

Rypel, A. L., M. C. Miner, L. Hitt, A. Holmes, S. E. Baird, D. E. Cocherell, A. M. Schreier, and N. A. Fangue. 2022. “Origin and Abundance of Chinook Salmon in Putah Creek.” Annual report to the Solano County Water Agency 2021/2022.

Rypel, A. L., P. Saffarinia, C. C. Vaughn, L. Nesper, K. O’Reilly, C. A. Parisek, M. L. Miller, et al. 2021. “Goodbye to ‘Rough Fish’: Paradigm Shift in the Conservation of Native Fishes.” Fisheries 46: 605–616. 10.1002/fsh.10660.

Sass, G. G., A. L. Rypel, and J. D. Stafford. 2017. “Inland Fisheries Habitat Management: Lessons Learned from Wildlife Ecology and a Proposal for Change.” Fisheries 42: 197–209. 10.1080/03632415.2017.1276344.

Schnell, I. B., K. Bohmann, and M. T. P. Gilbert. 2015. “Tag Jumps Illuminated – Reducing Sequence-to-Sample Misidentifications in Metabarcoding Studies.” Molecular Ecology Resources 15: 1289–1303. 10.1111/1755-0998.12402.

Seymour, M., F. K. Edwards, B. J. Cosby, I. Bista, P. M. Scarlett, F. L. Brailsford, H. C. Glanville, M. de Bruyn, G. R. Carvalho, and S. Creer. 2021. “Environmental DNA Provides Higher Resolution Assessment of Riverine Biodiversity and Ecosystem Function via Spatio-Temporal Nestedness and Turnover Partitioning.” Communications Biology 4: 512. 10.1038/s42003-021-02031-2.

Skelton, J., A. Cauvin, and M. E. Hunter. 2023. “Environmental DNA Metabarcoding Read Numbers and Their Variability Predict Species Abundance, but Weakly in Non Dominant Species.” Environmental DNA 5: 1092–1104. 10.1002/edn3.355.

Strayer, D. L., and D. Dudgeon. 2010. “Freshwater Biodiversity Conservation: Recent Progress and Future Challenges.” Journal of the North American Benthological Society 29: 344–358. 10.1899/08-171.1.

Thompson, K. 1960. “Historic Flooding in the Sacramento Valley.” Pacific Historical Review 29: 349–360. 10.2307/3636308.

Tonkin, J. D., M. T. Bogan, N. Bonada, B. Rios-Touma, and D. A. Lytle. 2017. “Seasonality and Predictability Shape Temporal Species Diversity.” Ecology 98: 1201–1216. 10.1002/ecy.1761.

Tonkin, J. D., J. D. Olden, D. M. Merritt, L. V. Reynolds, J. S. Rogosch, and D. A. Lytle. 2021. “Designing Flow Regimes to Support Entire River Ecosystems.” Frontiers in Ecology and the Environment 19: 326–333. 10.1002/fee.2348.

Vanni, M. J. 2002. “Nutrient Cycling by Animals in Freshwater Ecosystems.” Annual Review of Ecology and Systematics 33: 341–370. 10.1146/annurev.ecolsys.33.010802.150519.

Vannote, R. L., G. W. Minshall, K. W. Cummins, J. R. Sedell, and C. E. Cushing. 1980. “The River Continuum Concept.” Canadian Journal of Fisheries and Aquatic Sciences 37: 130–137. 10.1139/f80-017.

Walter, R. P., E. S. Gnyra, L. I. Söderberg, and D. D. Heath. 2014. “Rapid Genetic Identification of Brown Bullhead (*Ameiurus nebulosus*), Black Bullhead (*Ameiurus melas*) and Their Hybrids.” Conservation Genetics Resources 6: 507–509. 10.1007/s12686-014-0161-4.

Wang, Q., G. M. Garrity, J. M. Tiedje, and J. R. Cole. 2007. “Naïve Bayesian Classifier for Rapid Assignment of rRNA Sequences into the New Bacterial Taxonomy.” Applied and Environmental Microbiology 73: 5261–5267. 10.1128/AEM.00062-07.

Ward, D. L., A. F. Casper, T. D. Counihan, J. M. Bayer, I. R. Waite, J. J. Kosovich, C. G. Chapman, et al. 2017. “Long-Term Fish Monitoring in Large Rivers: Utility of ‘Benchmarking’ across Basins.” Fisheries 42: 100–114. 10.1080/03632415.2017.1276330.

Watson, J. M., S. M. Coghlan Jr, J. Zydlewski, D. B. Hayes, and I. A. Kiraly. 2018. “Dam Removal and Fish Passage Improvement Influence Fish Assemblages in the Penobscot River, Maine.” Transactions of the American Fisheries Society 147: 525–540. 10.1002/tafs.10053.

Wilsey, B. J., D. R. Chalcraft, C. M. Bowles, and M. R. Willig. 2005. “Relationships among Indices Suggest That Richness Is an Incomplete Surrogate for Grassland Biodiversity.” Ecology 86: 1178–1184. 10.1890/04-0394.

Wright, E. S. 2024. “Fast and Flexible Search for Homologous Biological Sequences with DECIPHER v3.” The R Journal 16: 191–200.

Yates, M. C., A. Van Nynatten, I. Smith, E. Spice, J. Leblanc, B. Thalinger, N.E. Mandrak, R. Bajno, C. M. Boston, H. MacIsaac, and D. Heath. 2025. “Longer Amplicon Metabarcoding Primers Enhance Fish Taxonomic Resolution in Environmental DNA Samples.” Canadian Journal of Fisheries and Aquatic Sciences 82: 1–14. 10.1139/cjfas-2025-0211.

Yarnell, S. M., G. E. Petts, J. C. Schmidt, A. A. Whipple, E. E. Beller, C. N. Dahm, P. Goodwin, and J. H. Viers. 2015. “Functional Flows in Modified Riverscapes: Hydrographs, Habitats and Opportunities.” BioScience 65: 963–972. 10.1093/biosci/biv102.

Yarnell, S. M., E. D. Stein, J. A. Webb, T. Grantham, R. A. Lusardi, J. Zimmerman, R. A. Peek, B. A. Lane, J. Howard, and S. Sandoval Solis. 2020. “A Functional Flows Approach to Selecting Ecologically Relevant Flow Metrics for Environmental Flow Applications.” River Research and Applications 36: 318–324. 10.1002/rra.3575.

Yeager, L. A., C. A. Layman, and J. E. Allgeier. 2011. “Effects of Habitat Heterogeneity at Multiple Spatial Scales on Fish Community Assembly.” Oecologia 167: 157–168. 10.1007/s00442-011-1959-3.

Young, R. T., J. C. Boase, J. T. Buszkiewicz, J. C. Dean, and J. T. McCarter. 2023. “Field Evaluation of Electrofishing Response Thresholds for Adult Grass Carp (*Ctenopharyngodon idella*).” North American Journal of Fisheries Management 43(3): 859–868. 10.1002/nafm.10899.

Yu, G., D. K. Smith, H. Zhu, Y. Guan, and T. T. Y. Lam. 2017. “GGTREE: An R Package for Visualization and Annotation of Phylogenetic Trees with Their Covariates and Other Associated Data.” Methods in Ecology and Evolution 8(1): 28–36. 10.1111/2041-210X.12628.

Zimmerman, J. K., D. M. Carlisle, J. T. May, K. R. Klausmeyer, T. E. Grantham, L. R. Brown, and J. K. Howard. 2018. “Patterns and Magnitude of Flow Alteration in California, USA.” Freshwater Biology 63(8): 859–873. 10.1111/fwb.13058.

