## Supplementary Material for "Community assembly dynamics in a regulated stream: Insights from environmental DNA"

**Table S1.** Sampling events for eDNA collection in Putah Creek (California, USA) from 2019 to 2021. Each row corresponds to a unique sampling event and includes the date, site name, and whether a concurrent conventional fish survey was conducted. This table supports interpretation of the bar plot in Supplementary Figure S2 by indicating the sampling event number. Additional site metadata is available in Table 1 in the main text.

| Sampling event | Date | Site | Concurrent conventional survey |
|---:|---|---|---|
| 1 | 19-DEC-2019 | Stebbins Cold Canyon | NA |
| 2 | 19-DEC-2019 | Step Weir 1 | NA |
| 3 | 19-DEC-2019 | Mace Boulevard, downstream | NA |
| 4 | 25-MAY-2020 | Russell Ranch | Fyke trap |
| 5 | 28-MAY-2020 | Stebbins Cold Canyon | NA |
| 6 | 28-MAY-2020 | Step Weir 1 | NA |
| 7 | 28-MAY-2020 | Mace Boulevard, downstream | NA |
| 8 | 19-OCT-2020 | Mace Boulevard, upstream | Electrofishing |
| 9 | 19-OCT-2020 | Old Davis Road | Electrofishing |
| 10 | 19-OCT-2020 | Pedrick Road | Electrofishing |
| 11 | 17-NOV-2020 | Russell Ranch | NA |
| 12 | 12-APR-2021 | Step Weir 2 | Rotary Screw Trap |
| 13 | 12-APR-2021 | Stebbins Cold Canyon | NA |
| 14 | 12-APR-2021 | Mace Boulevard, upstream | NA |
| 15 | 27-MAY-2021 | Step Weir 2 | Rotary Screw Trap |
| 16 | 18-OCT-2021 | Mace Boulevard, upstream | Electrofishing |
| 17 | 19-OCT-2021 | Old Davis Road | NA |
| 18 | 19-OCT-2021 | Pedrick Road | Electrofishing |
| 19 | 20-OCT-2021 | Interstate 505 Crossing | Electrofishing |
| 20 | 20-OCT-2021 | Winters Putah Creek Park | Electrofishing |
| 21 | 30-OCT-2021 | Fire Ring Picnic Grounds | Multiple methods |

**Table S2.** MiFish-U primer sequences (bolded; Miya et al. 2015) with "heterogeneity" spacers (underlined; Fadrosh et al. 2014) and the Illumina adapter overhang sequence were pooled in equimolar amounts for the amplicon PCR step of metabarcoding library preparation.

| Primer name | Spacer | Primer sequence (5'-3') |
|---|---|---|
| MiFish-U-F | 0 bp | TCGTCGGCAGCGTCAGATGTGTATAAGAGACAG**GTCGGTAAAACTCGTGCCAGC** |
| | 2 bp | TCGTCGGCAGCGTCAGATGTGTATAAGAGACAGTA**GTCGGTAAAACTCGTGCCAGC** |
| | 4 bp | TCGTCGGCAGCGTCAGATGTGTATAAGAGACAGCATA**GTCGGTAAAACTCGTGCCAGC** |
| | 6 bp | TCGTCGGCAGCGTCAGATGTGTATAAGAGACAGCGCATA**GTCGGTAAAACTCGTGCCAGC** |
| MiFish-U-R | 0 bp | GTCTCGTGGGCTCGGAGATGTGTATAAGAGACAG**CATAGTGGGGTATCTAATCCCAGTTTG** |
| | 2 bp | GTCTCGTGGGCTCGGAGATGTGTATAAGAGACAGTA**CATAGTGGGGTATCTAATCCCAGTTTG** |
| | 4 bp | GTCTCGTGGGCTCGGAGATGTGTATAAGAGACAGCATA**CATAGTGGGGTATCTAATCCCAGTTTG** |
| | 6 bp | GTCTCGTGGGCTCGGAGATGTGTATAAGAGACAGCGCATA**CATAGTGGGGTATCTAATCCCAGTTTG** |

**Table S3.** Amplicon PCR reagent volumes and concentrations.

| Component | Volume per reaction (µl) | Final concentration |
|---|---|---|
| KAPA HiFi Fidelity Buffer (5x) | 5 | 1x |
| dNTP mix (10 mM each dNTP) | 0.75 | 300 µM each dNTP |
| KAPA HiFi HotStart DNA Polymerase | 5 | 1 U |
| MiFish-U-F primer | 0.75 | 0.3 µM |
| MiFish-U-R primer | 0.75 | 0.3 µM |
| PCR-grade water | 11.75 | - |
| DNA template | 4 | - |
| | **25 µl total volume** | |

**Table S4.** Thermocycling conditions for the amplicon PCR step of metabarcoding library preparation.

| Step | Time | Temperature | Cycles |
|---|---|---|---|
| Initial denature | 3:00 | 95ºC | 1 |
| Denature | 0:15 | 98ºC | |
| Anneal | 0:20 | 68ºC | 35 |
| Extension | 0:15 | 72ºC | |
| Extension | 5:00 | 72ºC | 1 |
| Hold | Hold | 12ºC | 1 |

**Table S5.** Final fish read counts after quality-control and decontamination of Illumina sequencing data.

| SampleID | ReadCount |
|---|---|
| PC_001 | 64,658 |
| PC_002 | 75,985 |
| PC_003 | 62,614 |
| PC_004 | 83,779 |
| PC_005 | 89,144 |
| PC_006 | 31,821 |
| PC_007 | 78,502 |
| PC_008 | 74,373 |
| PC_009 | 45,260 |
| PC_018 | 59,598 |
| PC_019 | 51,567 |
| PC_020 | 63,617 |
| PC_025 | 61,009 |
| PC_026 | 59,674 |
| PC_027 | 59,813 |
| PC_028 | 34,134 |
| PC_029 | 72,600 |
| PC_030 | 52,672 |
| PC_031 | 37,849 |
| PC_035 | 44,533 |
| PC_036 | 6,112 |
| PC_037 | 18,952 |
| PC_038 | 55,728 |
| PC_039 | 67,939 |
| PC_040 | 96,045 |
| PC_041 | 78,111 |
| PC_043 | 5,733 |
| PC_044 | 75,107 |
| PC_045 | 81,737 |
| PC_049 | 11,708 |

| SampleID | ReadCount |
|---|---|
| PC_051 | 29,690 |
| PC_052 | 61,444 |
| PC_053 | 38,129 |
| PC_055 | 15,370 |
| PC_056 | 17,720 |
| PC_057 | 6,841 |
| PC_058 | 11,292 |
| PC_059 | 7,531 |
| PC_061 | 2,772 |
| PC_062 | 21,036 |
| PC_063 | 25,762 |
| PC_070 | 75,329 |
| PC_071 | 86,603 |
| PC_072 | 78,307 |
| PC_074 | 67,352 |
| PC_075 | 78,436 |
| PC_076 | 101,818 |
| PC_078 | 52,601 |
| PC_079 | 103,113 |
| PC_080 | 96,865 |
| PC_082 | 31,102 |
| PC_083 | 36,207 |
| PC_084 | 57,495 |
| PC_085 | 45,459 |
| PC_086 | 49,431 |
| PC_087 | 41,710 |
| PC_090 | 38,108 |
| PC_091 | 54,506 |
| PC_092 | 52,398 |

**Table S6.** Reference sequence species pairwise genetic distances <3% at the MiFish barcode region.

| Species1 | Species2 | Genetic distance | Percent similarity |
|---|---|---|---|
| common carp | goldfish | 0.0231 | 97.69 |
| hitch | California roach | 0.0114 | 98.86 |
| Sacramento pikeminnow | California roach | 0.0286 | 97.14 |
| Sacramento pikeminnow | hitch | 0.0286 | 97.14 |
| kokanee | rainbow trout | 0.0059 | 99.41 |
| Chinook salmon | rainbow trout | 0.0235 | 97.65 |
| Chinook salmon | kokanee | 0.0176 | 98.24 |

**Table S7.** Site-level ecological metrics pooled by date for Putah Creek eDNA samples. Species richness was calculated for all fishes (A), native fishes (N) and non-native fishes (NN). Shannon diversity and evenness (Pielou's J) should be interpreted with caution due to potential taxonomic biases inherent to DNA amplification (i.e., PCR bias).

| Site | Sampling event | Season | Species richness | | | Shannon diversity | Evenness |
|---|---|---|---|---|---|---|---|
| | | | A | N | NN | | |
| Mace Boulevard, downstream | 3 | Fall | 20 | 7 | 13 | 2.30 | 0.77 |
| | 7 | Spring | 14 | 3 | 11 | 1.76 | 0.67 |
| Mace Boulevard, upstream | 8 | Fall | 14 | 3 | 11 | 1.37 | 0.52 |
| | 14 | Spring | 14 | 4 | 10 | 1.96 | 0.74 |
| | 16 | Fall | 19 | 4 | 15 | 2.00 | 0.68 |
| Old Davis Road | 9 | Fall | 10 | 3 | 7 | 1.69 | 0.74 |
| | 17 | Fall | 19 | 5 | 14 | 2.32 | 0.79 |
| Fire Ring Picnic Grounds | 21 | Fall | 15 | 6 | 9 | 1.86 | 0.69 |
| Pedrick Road | 10 | Fall | 9 | 4 | 5 | 1.37 | 0.62 |
| | 18 | Fall | 10 | 5 | 5 | 1.42 | 0.61 |
| Russell Ranch | 4 | Spring | 12 | 5 | 7 | 1.21 | 0.48 |
| | 11 | Fall | 8 | 5 | 3 | 1.66 | 0.80 |
| Step Weir 1 | 2 | Fall | 11 | 7 | 4 | 1.71 | 0.71 |
| | 6 | Spring | 11 | 7 | 4 | 1.69 | 0.70 |
| Step Weir 2 | 12 | Spring | 10 | 8 | 2 | 1.34 | 0.58 |
| | 15 | Spring | 15 | 8 | 7 | 1.17 | 0.43 |
| Interstate 505 Crossing | 19 | Fall | 11 | 7 | 4 | 1.54 | 0.64 |
| Winters Putah Creek Park | 20 | Fall | 12 | 7 | 5 | 1.20 | 0.48 |
| Stebbins Cold Canyon | 1 | Fall | 11[1] | 6 | 5[1] | 1.45 | 0.60 |
| | 5 | Spring | 8 | 6 | 2 | 1.19 | 0.57 |
| | 13 | Spring | 8 | 5 | 3 | 1.14 | 0.55 |

[1]Detection of two non-native species not known from Putah Creek may reflect eDNA transport from Lake Berryessa, just upstream from this site.

**Table S8.** Exploratory mean rank shift and species turnover calculations across years and seasons for eDNA sampling sites with multiple time points. For season-to-season comparisons at river kilometer (km) 11.1, Supplementary Figure S6 displays the mean value.

| river_km | sampling_event_1 | sampling_event_2 | comparison_type | category | MRS | MRS_avg_for_plot | turnover | turnover_avg_for_plot |
|---|---|---|---|---|---|---|---|---|
| 10.6 | 3 | 7 | Fall 2019 → Spring 2020 | Season to Season | 2.15 | 2.15 | 0.38 | 0.38 |
| 34.4 | 2 | 6 | Fall 2019 → Spring 2020 | Season to Season | 2.20 | 2.20 | 0.17 | 0.17 |
| 51 | 1 | 5 | Fall 2019 → Spring 2020 | Season to Season | 0.40 | 0.40 | 0.64 | 0.64 |
| 51 | 5 | 13 | Spring 2020 → Spring 2021 | Year to Year | 0.40 | 0.40 | 0.55 | 0.55 |
| 11.1 | 8 | 16 | Fall 2020 → Fall 2021 | Year to Year | 1.85 | 1.85 | 0.35 | 0.35 |
| 16.9 | 9 | 17 | Fall 2020 → Fall 2021 | Year to Year | 1.80 | 1.80 | 0.47 | 0.47 |
| 20.8 | 10 | 18 | Fall 2020 → Fall 2021 | Year to Year | 0.75 | 0.75 | 0.27 | 0.27 |
| 11.1 | 8 | 14 | Fall 2020 → Spring 2021 | Season to Season | 2.73 | 2.59 | 0.35 | 0.35 |
| 11.1 | 14 | 16 | Spring 2021 → Fall 2021 | Season to Season | 2.46 | 2.59 | 0.35 | 0.35 |

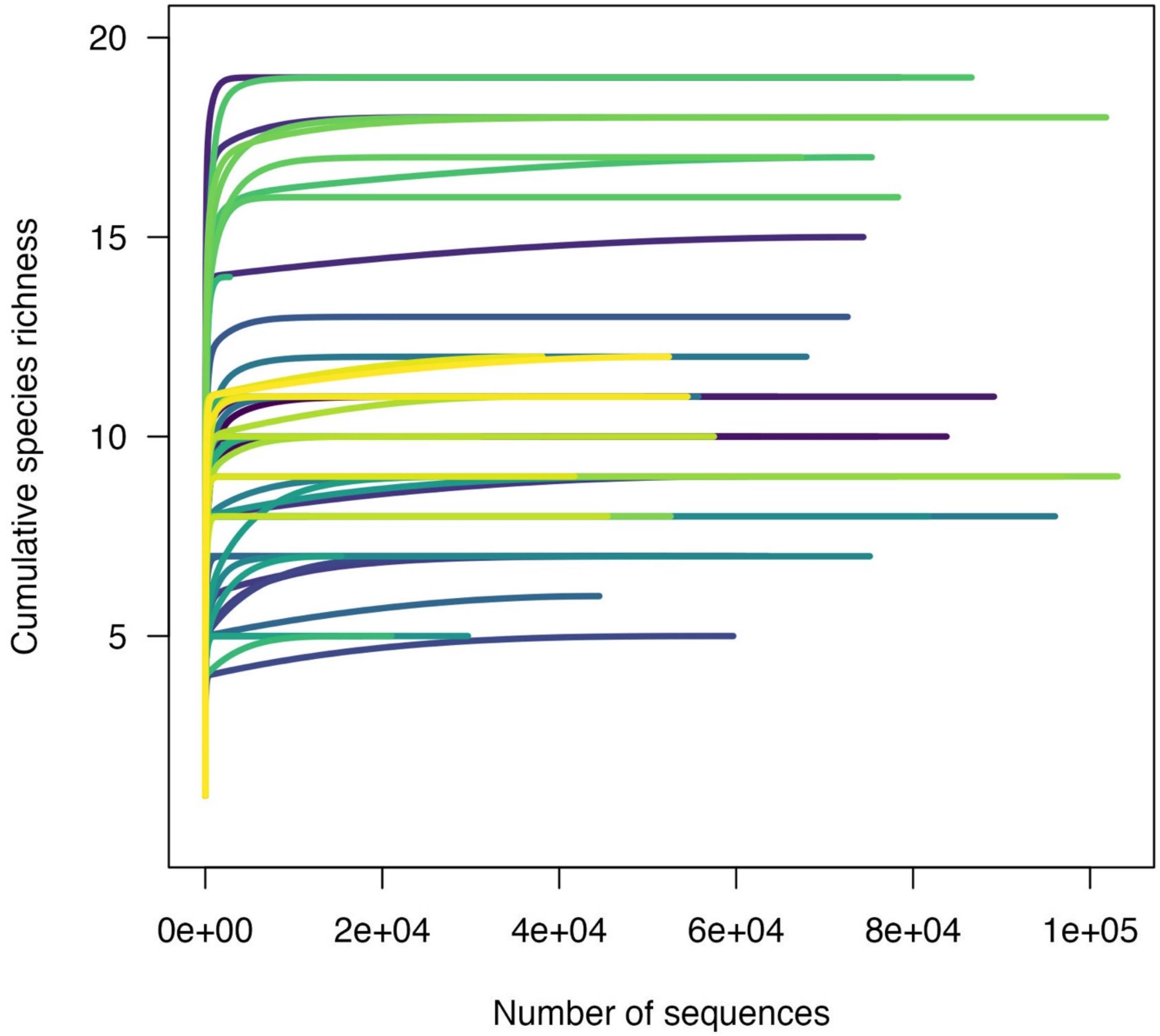

**Figure S1.** Species accumulation curves where each line represents the cumulative number of unique fish species detected as sequencing depth increases. Asymptotic curves indicate that sequencing depth was sufficient to characterize the species richness of samples.

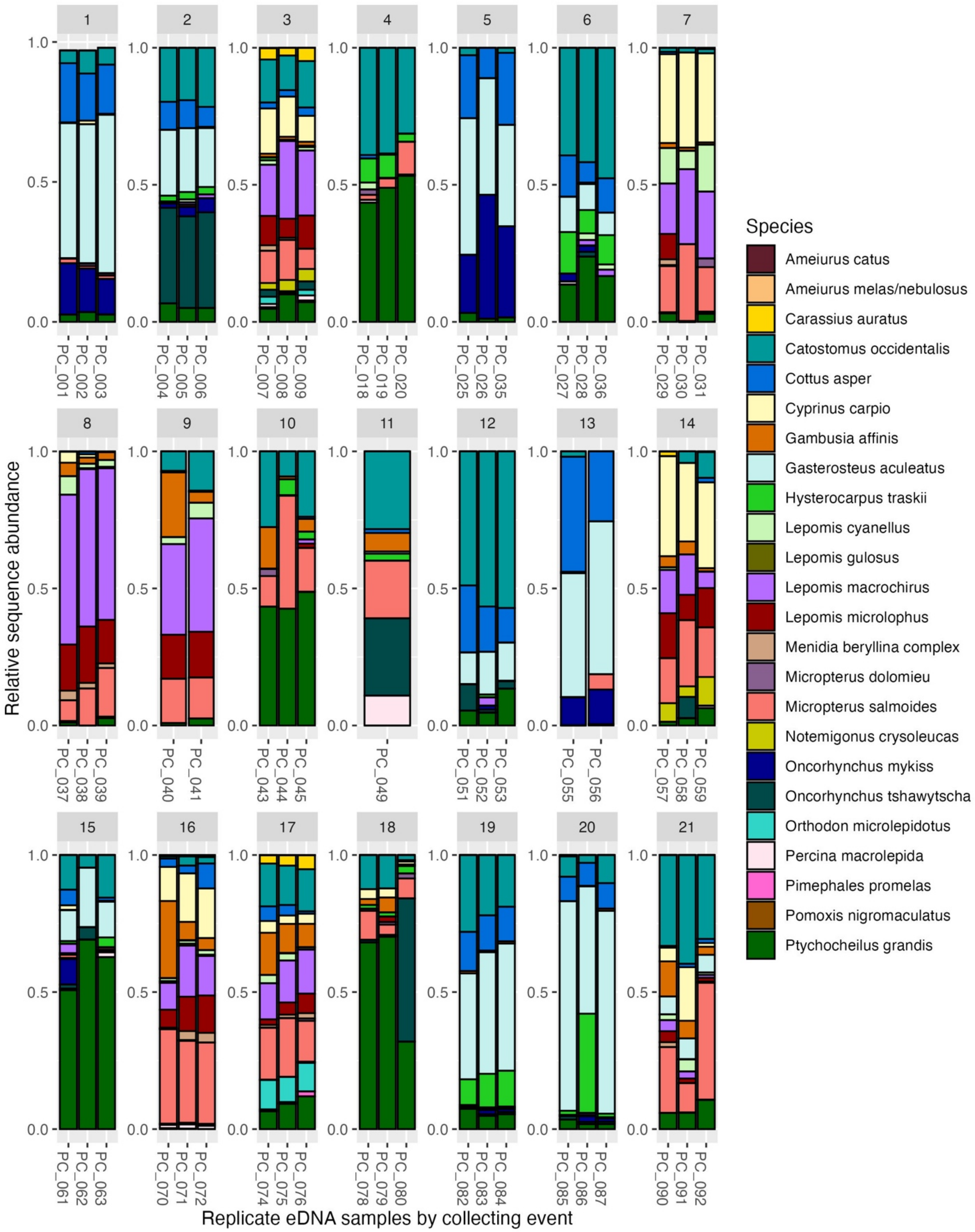

**Figure S2.** Relative read abundance of fish species detected in eDNA samples collected from Putah Creek between 2019 and 2021. Each bar represents a single eDNA replicate, grouped by sampling event (see Supplementary Table S1 for sampling event information); three sampling events had fewer than three replicates due to amplification failure. Kokanee and

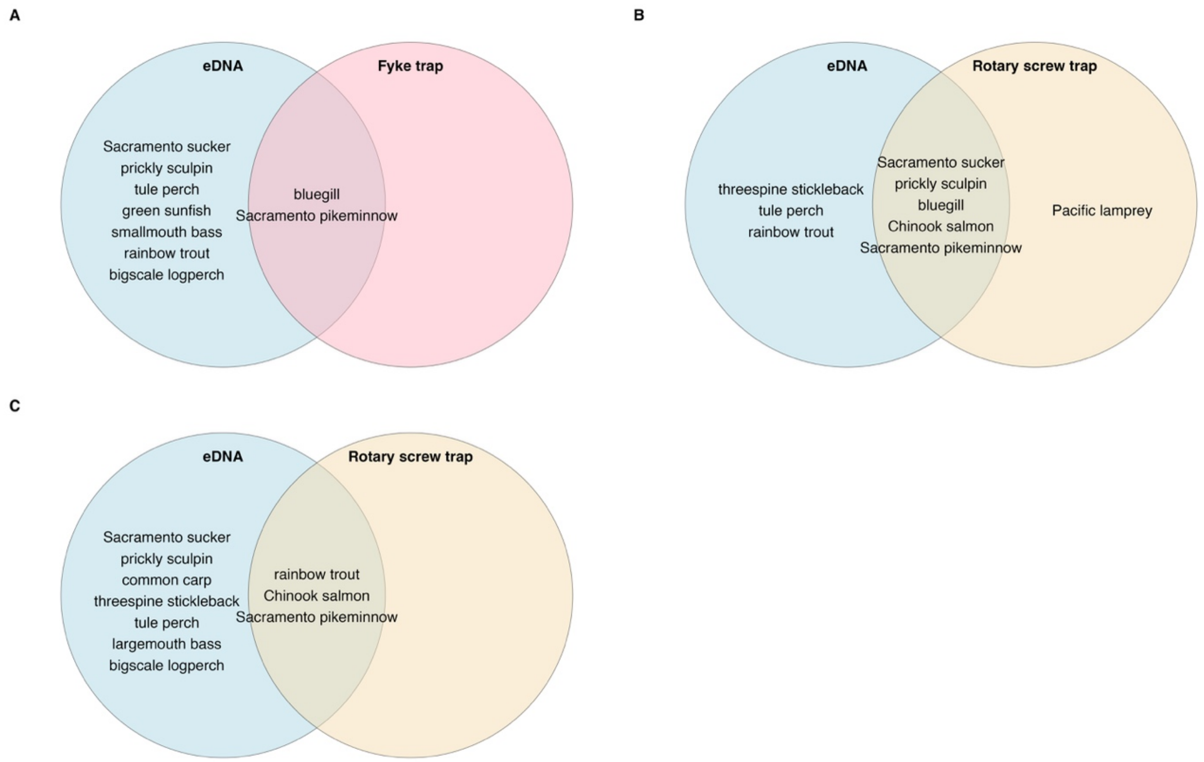

**Figure S3.** Venn diagrams comparing fish species detected by eDNA and trap sampling: fyke trap at RRAN in spring 2020 (A) and rotary screw trap at SW1D in spring 2021 (B, April; C, May).

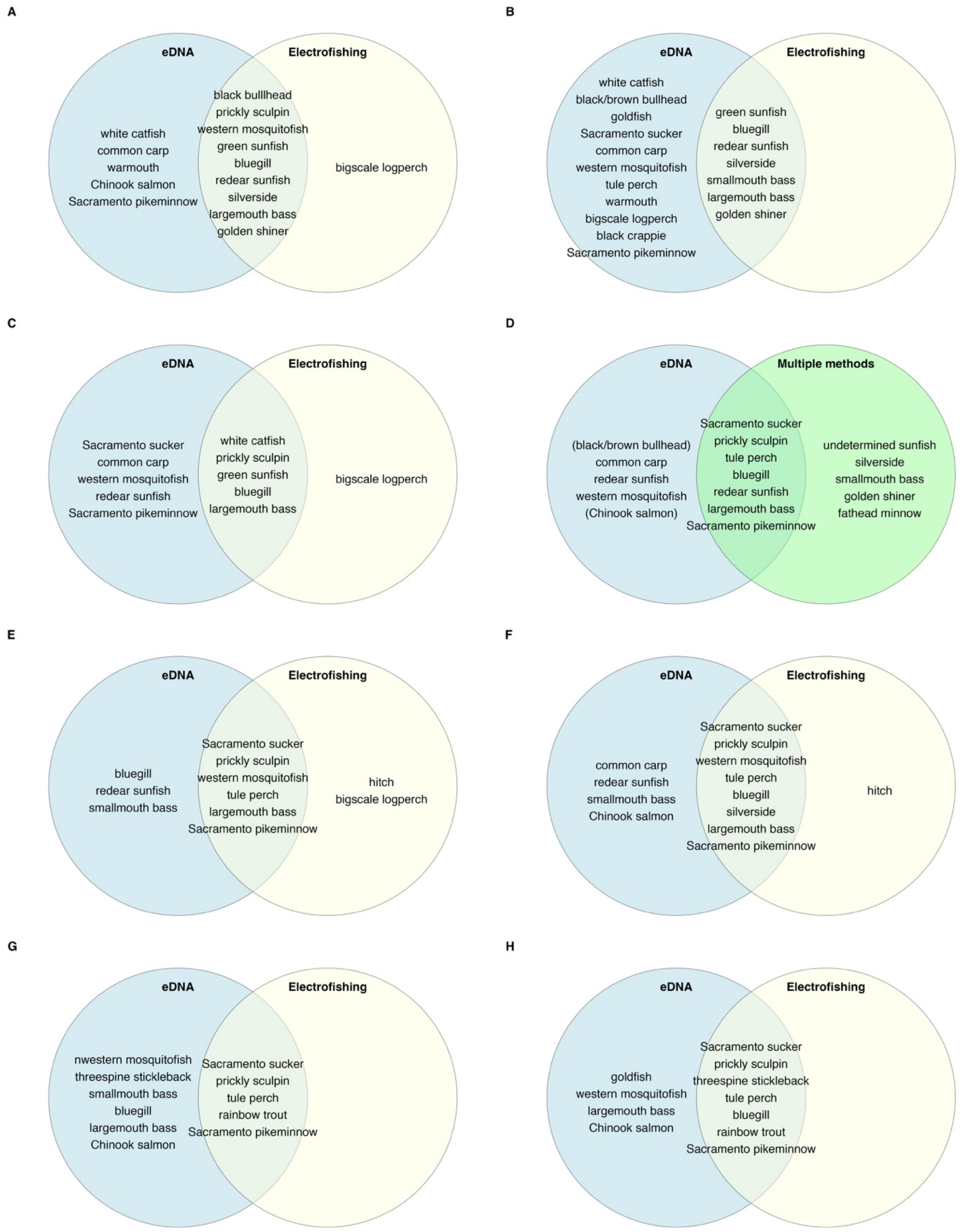

**Figure S4.** Venn diagrams comparing fish species detected by eDNA and electrofishing. Sampling event D used electrofishing, seines, and traps.

**A**

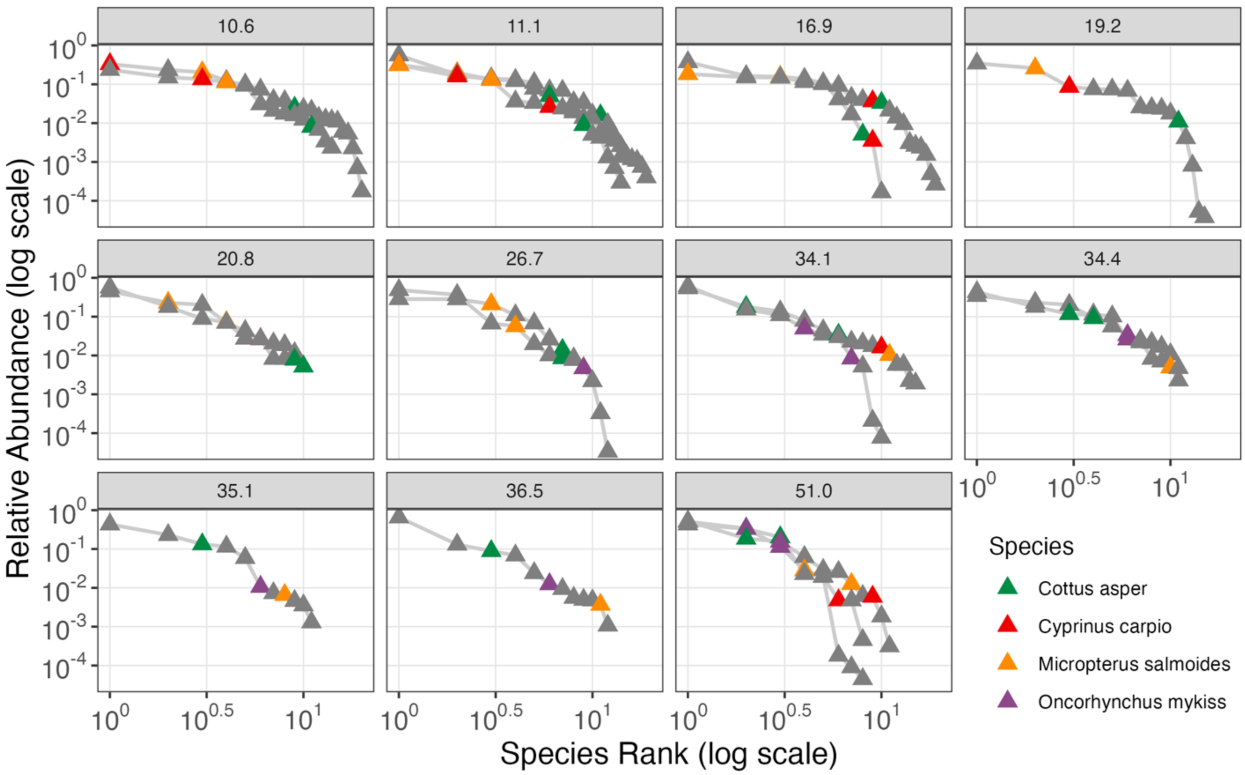
10.6
11.1
16.9
19.2
20.8
26.7
34.1
34.4
35.1
36.5
51.0
Relative Abundance (log scale)
Species Rank (log scale)
Species
Cottus asper
Cyprinus carpio
Micropterus salmoides
Oncorhynchus mykiss

B

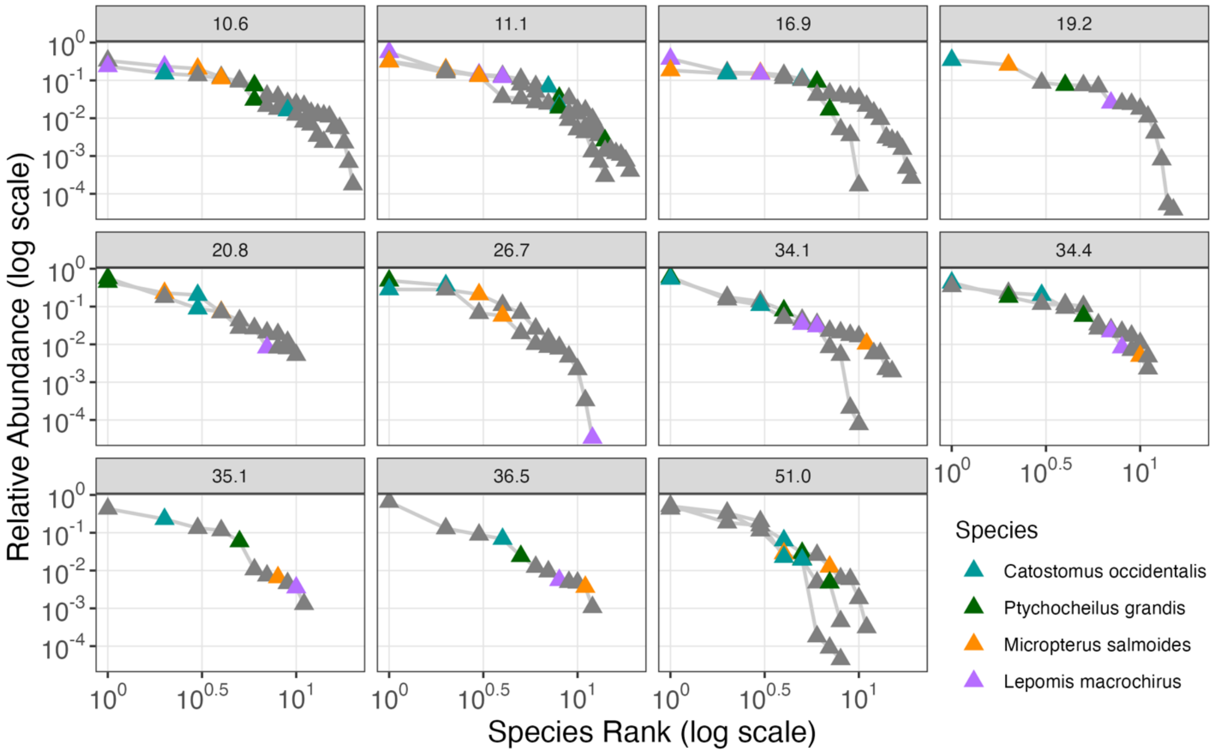

**Figure S5.** Rank-abundance curves (Whittaker plots) highlighting focal species detected using eDNA in Putah Creek arranged by river kilometer (km) site. Each curve represents pooled replicates from one sampling event (n = 1 to 3 sampling events/site) and flatter slopes indicate higher community evenness. Species are ranked by relative abundance on a log-log scale. Colored points indicate (**A**) species highlighted in Jacinto et al. (2023) derived from historical electrofishing data (1993-2017): native *Cottus asper* (prickly sculpin) and *Oncorhynchus mykiss* (rainbow trout), and non-native *Cyprinus carpio* (common carp) and *Micropterus salmoides* (largemouth bass) or (**B**) the four most abundant species in the eDNA dataset by relative read abundance: native *Catostomus occidentalis* (Sacramento sucker) and *Ptychocheilus grandis* (Sacramento pikeminnow), and non-native *Micropterus salmoides* (largemouth bass) and *Lepomis macrochirus* (bluegill); all other species are shown in gray.

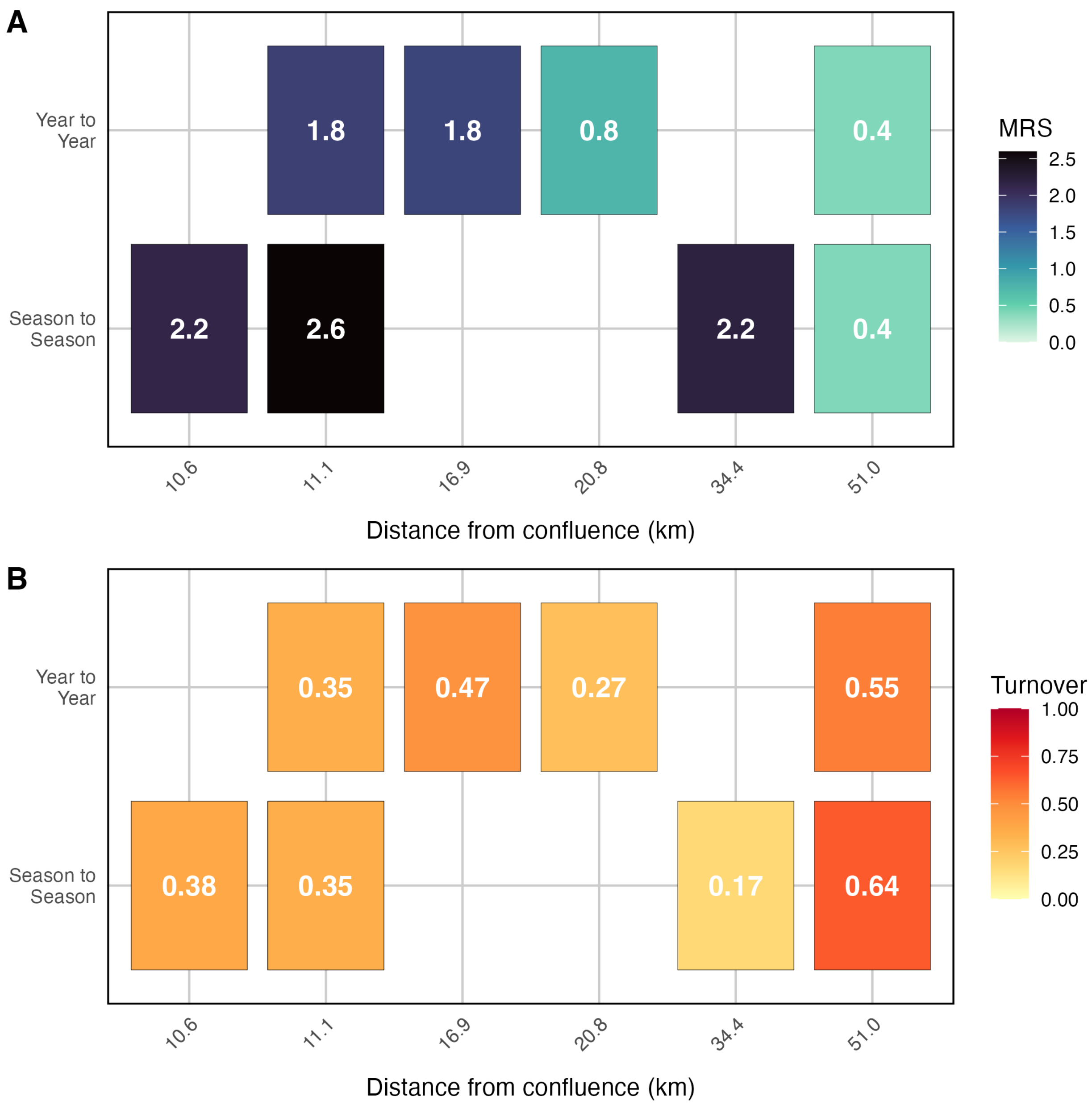

**Figure S6.** Exploratory mean rank shift and species turnover visualized from sequential sampling events (Supplementary Table S6); values should be interpreted with caution as these metrics are typically applied to long-term datasets with annual sampling intervals.

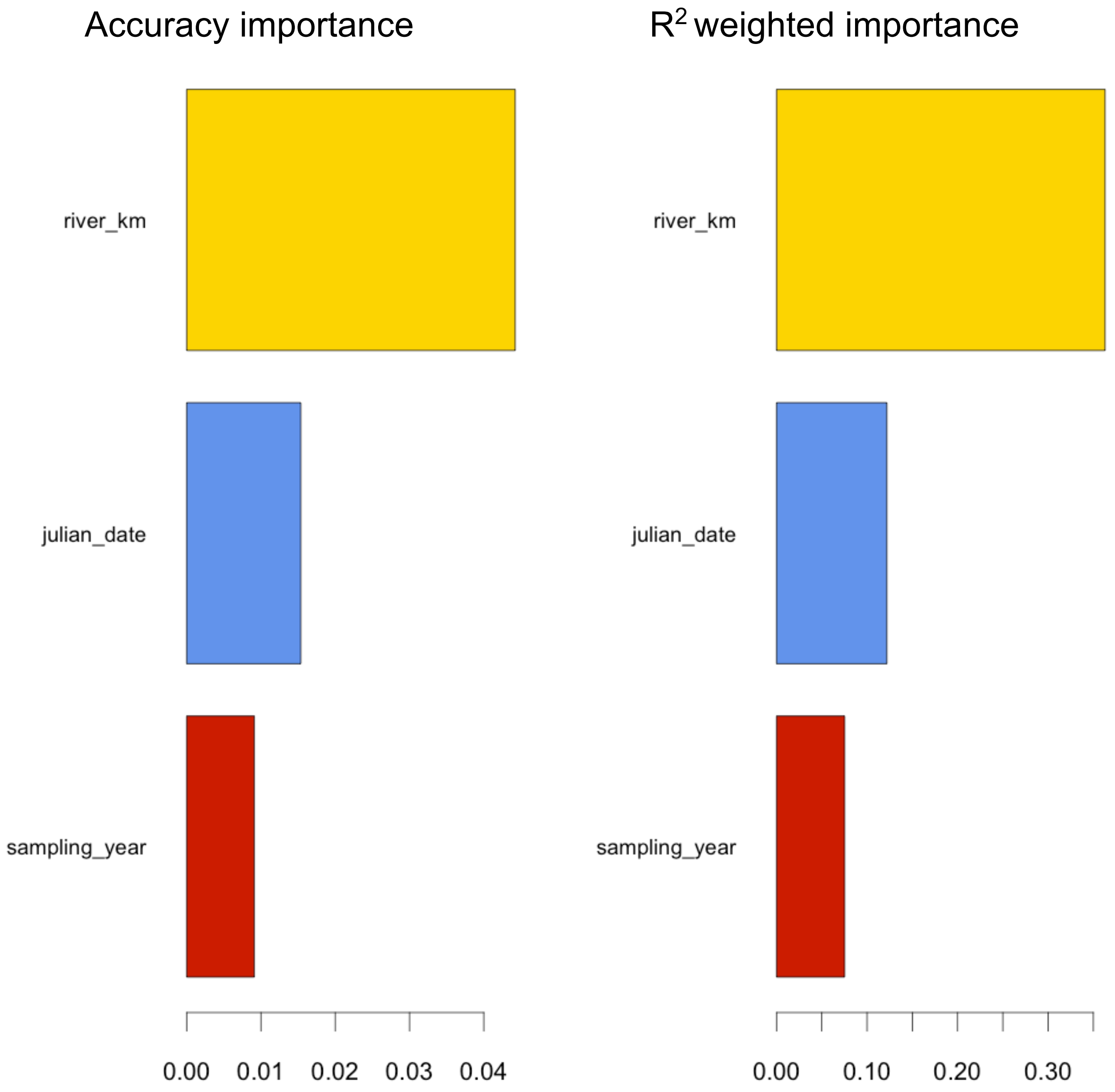

**Figure S7.** Variable importance in the gradient forest model detected in Putah Creek, showing each predictor's contribution to species turnover. River kilometer (a proxy for distance upstream from the confluence of Putah Creek with the Yolo Bypass) had the highest accuracy importance (left) and $R^2$-weighted importance (right), compared with Julian date and year. Species-level importance in the gradient forest model is reported in the main text (Figure 7).
